# STAG2 promotes the myelination transcriptional program in oligodendrocytes

**DOI:** 10.1101/2021.10.10.463866

**Authors:** Ningyan Cheng, Mohammed Kanchwala, Bret M. Evers, Chao Xing, Hongtao Yu

**Affiliations:** Department of Pharmacology, University of Texas Southwestern Medical Center, Dallas, TX 75390, USA; Eugene McDermott Center for Human Growth and Development, University of Texas Southwestern Medical Center, Dallas, TX 75390, USA; Division of Neuropathology, University of Texas Southwestern Medical Center, Dallas, TX, 75390, USA; Department of Bioinformatics, Department of Population and Data Sciences, University of Texas Southwestern Medical Center, Dallas, TX 75390, USA; Westlake Laboratory of Life Sciences and Biomedicine, Hangzhou, China; School of Life Sciences, Westlake University, Hangzhou, China; Institute of Biology, Westlake Institute for Advanced Study, Hangzhou, China

## Abstract

Cohesin folds chromosomes via DNA loop extrusion. Cohesin-mediated chromosome loops regulate transcription by shaping long-range enhancer-promoter interactions, among other mechanisms. Mutations of cohesin subunits and regulators cause human developmental diseases termed cohesinopathy. Vertebrate cohesin consists of SMC1, SMC3, RAD21, and either STAG1 or STAG2. To probe the physiological functions of cohesin, we created conditional knockout (cKO) mice with *Stag2* deleted in the nervous system. *Stag2* cKO mice exhibit growth retardation, neurological defects, and premature death, in part due to insufficient myelination of nerve fibers. *Stag2* cKO oligodendrocytes exhibit delayed maturation and downregulation of myelination-related genes. *Stag2* loss reduces promoter-anchored loops at downregulated genes in oligodendrocytes. Thus, STAG2-cohesin generates promoter-anchored loops at myelination-promoting genes to facilitate their transcription. Our study implicates defective myelination as a contributing factor to cohesinopathy and establishes oligodendrocytes as a relevant cell type to explore the mechanisms by which cohesin regulates transcription.

## INTRODUCTION

Chromosomes in a single human diploid cell, if linearly stitched together, span a length of more than two meters. They need to be properly folded to be housed in the cell nucleus with a diameter of 10 µm. Chromosome folding occurs in a dynamic, structured way that regulates gene expression, and DNA replication and repair. Initially discovered as the molecular glue that tethers sister chromatids for segregation during mitosis (Haarhuis et al., 2014; Uhlmann, 2016; Yatskevich et al., 2019; Zheng and Yu, 2015), the cohesin complex has later been shown to be critical for structured chromosome folding and gene expression (Haarhuis et al., 2017; Rao et al., 2017; Schwarzer et al., 2017; Wutz et al., 2017).

Cohesin is loaded on chromosomes by the cohesin loader NIPBL. The cohesin–NIPBL complex can extrude DNA loops bi-directionally in an ATP-dependent manner (Davidson et al., 2019; Kim et al., 2019; Vian et al., 2018). The chromatin insulator CTCF has been proposed to block loop extrusion by cohesin, establishing topologically associated domains (TADs) and marking TAD boundaries. Chromatin interactions within each TAD are favored whereas inter-TAD interactions are disfavored. Thus, chromosome loops and TADs shape long-range cis-element interactions, such as promoter-enhancer interactions, thereby regulating transcription.

The vertebrate cohesin complex contains four core subunits: the SMC1–SMC3 heterodimeric ATPase, the kleisin subunit RAD21 that links the ATPase heads, and the HEAT-repeat protein STAG1 or STAG2. STAG1 and STAG2 bind to RAD21 in a mutually exclusive manner and create docking sites for several regulatory proteins, including CTCF (Hara et al., 2014; Li et al., 2020). STAG1 and STAG2 also interact with DNA and the SMC1–SMC3 hinge domains (Shi et al., 2020). STAG1 and STAG2 play redundant roles in sister-chromatid cohesion in cultured human cells, as both need to be simultaneously depleted to produce overt cohesion defects (Hara et al., 2014).

Mutations of NIPBL and cohesin subunits, including STAG2, result in human developmental diseases termed cohesinopathies, which affect multiple organs and systems (Remeseiro et al., 2013b; Soardi et al., 2017). In patients with cohesinopathies, mental retardation and neurological abnormalities caused by brain development defects are common (Piche et al., 2019). Dysregulation of gene transcription as a result of reduced cohesin functions has been suggested to underlie these developmental defects (De Koninck and Losada, 2016; Remeseiro et al., 2013a). In addition, several cohesin genes, including *STAG2*, are frequently mutated in a variety of human cancers (Martincorena and Campbell, 2015).

To better understand the physiological roles of cohesin, we deleted *Stag2* specifically in the nervous system in the mouse. The *Stag2* conditional knockout (cKO) mice exhibited deficient myelination. Loss of STAG2 delayed the maturation of oligodendrocytes and reduced chromosome loops in oligodendrocytes and impaired the transcription of myelination-related genes. Our findings establish the requirement for cohesin in proper gene expression in specific cell types and implicate defective myelination as a potential contributing factor to cohesinopathy.

## RESULTS

### *Stag2* ablation in the nervous system causes growth retardation and neurological defects

*Stag1* is required for mammalian embryonic development (Remeseiro et al., 2012), indicating that *Stag2* cannot compensate for the loss of *Stag1*. To examine the roles of *Stag2* during development in the mouse, we targeted a critical exon (exon 8) of *Stag2*, which is located on the X chromosome, using CRISPR-Cas9 (Figure S1A). The *Stag2^null^* embryos showed severe developmental defects and underwent necrosis by E11.5 days (Figure S1B). Thus, *Stag2* is required for mouse embryonic development, consistent with a recent report (De Koninck et al., 2020). *Stag1* and *Stag2* have non-redundant developmental functions.

Because whole-body knockout of *Stag2* caused embryonic lethality, we created a *Stag2* “floxed” mouse line (*Stag2^f/f^*) by homologous recombination with a template that contained two LoxP sites flanking exon 8 (Figure S2A,B). To study the physiological function of STAG2 in adult mice, we crossed the *Stag2^f/f^* mice with mice bearing the *Ert2-Cre* genomic insertion and generated *Stag2^f/y^;Ert2-Cre* progenies. The *Stag2^f/y^;Ert2-Cre* adult mice were injected with tamoxifen to induce *Stag2* deletion in the whole body (Figure S2C). Genotyping analysis of blood extracts showed that tamoxifen induced efficient disruption of the *Stag2* gene locus in *Stag2^f/y^;Ert2-Cre* mice (Figure S2D,E). These *Stag2*-deficient adult mice did not show early onset of spontaneous tumor formation, indicating that *Stag2* mutation alone in somatic cells of mice is insufficient to induce tumorigenesis. The *Stag2*-deficient mice also did not have other obvious adverse phenotypes (Figure S2F), except that they had slightly lower body weight (Figure S2G,H), probably due to tissue homeostasis alterations recently reported by others (De Koninck et al., 2020).

STAG2 mutations are found in human cohesinopathy patients with mental retardation and neuropsychiatric behaviors (Soardi et al., 2017). To study the function of STAG2 in the nervous system, we generated *Stag2* conditional knockout mice (*Stag2* cKO) by crossing *Stag2^f/f^* mice with *Nestin-Cre* mice (Giusti et al., 2014) (Figure 1A,B). The progenies were born in the Mendelian ratio, but *Stag2^f/y^;Cre* pups presented growth retardation and premature death (Figure 1C-E). More than 50% *Stag2^f/y^;Cre* mice died aged about 3 weeks while the rest died at about 4 months. *Stag2^f/y^* mice did not show differences discernible from WT littermates. Although *Stag2^f/y^;Cre* mice did not present microcephaly, they exhibited frequent hydrocephaly that might contribute to their premature death. The *Stag2^f/y^;Cre* mice displayed normal drinking and feeding behaviors (Figure 1F,G), but showed reduced plasma IGF-1 levels compared to the control mice (Figure 1H). *Stag2^f/y^;Cre* mice showed forepaw and hindlimb clasping (Figure 1I) and limb tremors (Movie 1), which were not seen in *Stag2^f/y^* mice. These data indicate that *Stag2* deficiency in the nervous system causes growth retardation and neurological defects.

**Figure 1.**
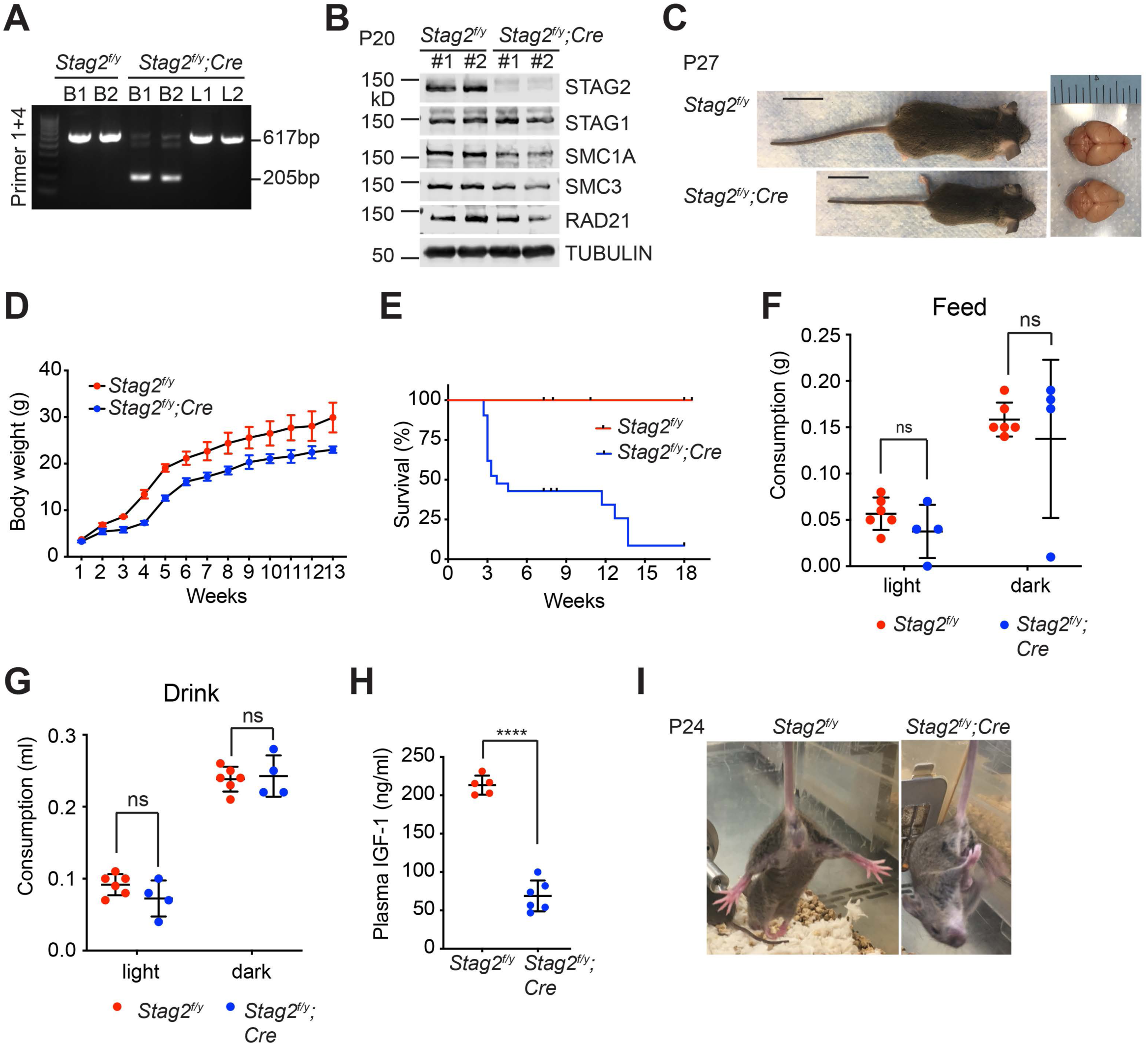
*Stag2* ablation in the mouse nervous system causes growth retardation and neurological defects. (A) PCR analysis of genomic DNA extracted from brains (BR) or livers (LV) of indicated mice. (B) Immunoblots of brain lysates of *Stag2^f/y^* and *Stag2^f/y^;Cre* mice. (C) Representative images of *Stag2^f/y^* and *Stag2^f/y^;Cre* mice. Scale bar = 2 cm. (D) Body weight of *Stag2^f/y^* and *Stag2^f/y^;Cre* mice at different age. Mean ± SD of at least three mice of the same age. (E) Survival curves of *Stag2^f/y^* (n = 12) and *Stag2^f/y^;Cre* (n = 21) mice. (F,G) Food (F) and water (G) consumption of 7- to 8-week-old *Stag2^f/y^* (n = 6) and *Stag2^f/y^;Cre* (n = 4) mice. Mean ± SD; ns, not significant. (H) Plasma IGF-1 levels of two-month-old *Stag2^f/y^* (n = 5) and *Stag2^f/y^;Cre* (n = 6) mice. Mean ± SD; ****p < 0.0001 (I) Representative images of limb-clasping responses of *Stag2^f/y^* and *Stag2^f/y^;Cre* mice.

### *Stag2* ablation causes hypomyelination

Hematoxylin and eosin staining of brain sections of *Stag2^f/y^;Cre* mice did not reveal overt anatomical defects (Figure S3A). To understand the origins of neurological defects caused by *Stag2* deletion, we analyzed the gene expression changes in *Stag2^f/y^;Cre* mouse brains by RNA-sequencing (RNA-seq) (Figure 2A). Compared with the control groups, 105 and 62 genes were significantly downregulated or upregulated by more than two folds, respectively, in the *Stag2*- deficient brains. The decreased expression of top differentially expressed genes (DEGs) was confirmed by reverse transcription quantitative PCR (RT-qPCR) (Figure 2B). Among the 105 downregulated DEGs in the brains of *Stag2* cKO mice, 44 were enriched in myelin (Figure 2C) (Thakurela et al., 2016). The Ingenuity Pathway Analysis (IPA) pinpoints cholesterol biosynthesis pathways as the most affected canonical pathways (Figure 2D). We further confirmed that the cholesterol biosynthesis precursors were reduced in *Stag2^f/y^;Cre* brains (Figure S3B).

**Figure 2.**
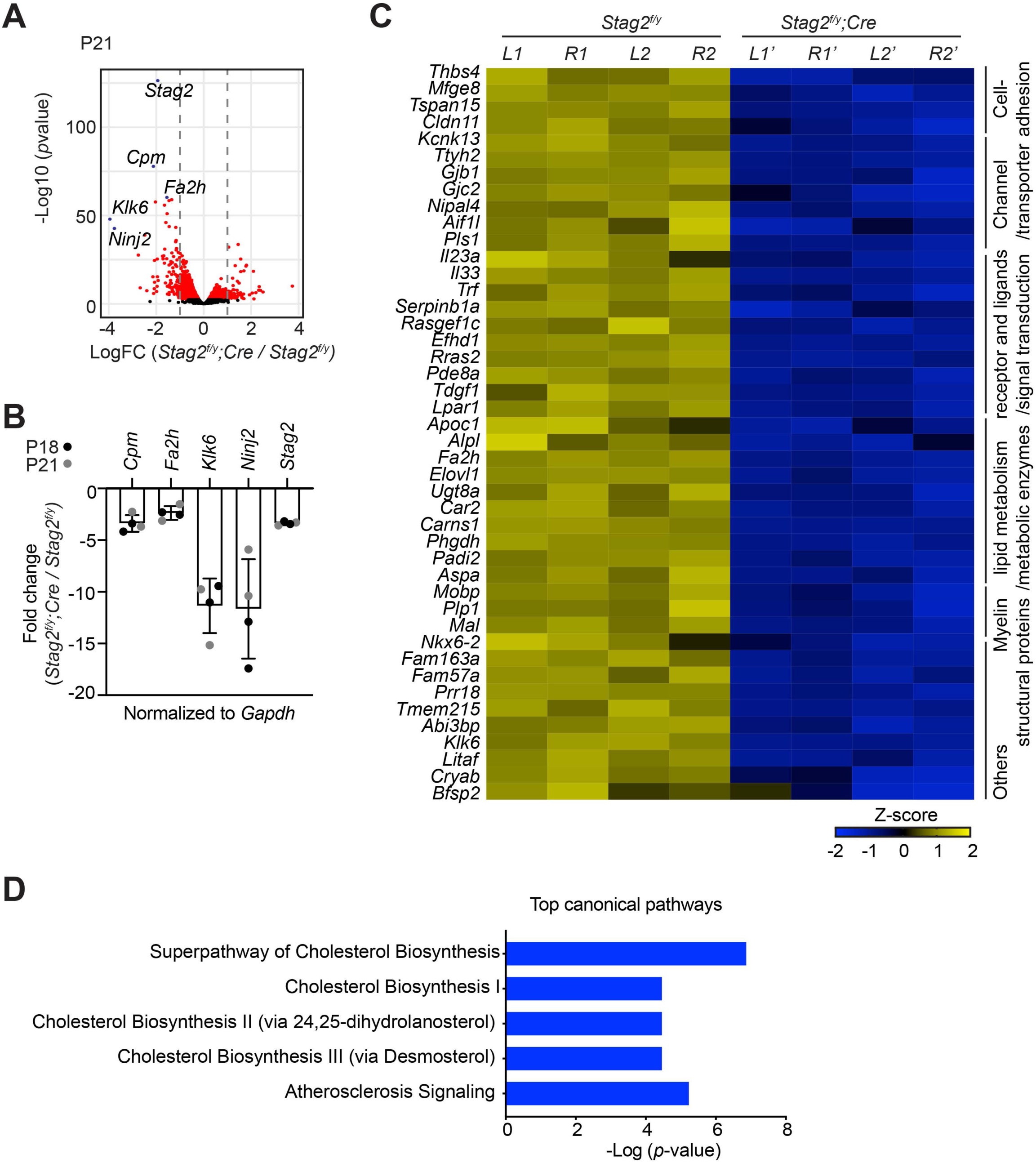
*Stag2* ablation in mouse brains downregulates the expression of myelin genes. (A) Volcano plot of bulk RNA sequencing results of *Stag2^f/y^* and *Stag2^f/y^;Cre* brain extracts. Top differentially expressed genes (DEGs) are colored blue and labeled. n = 4 pairs of P21 *Stag2^f/y^* and *Stag2^f/y^;Cre* brain hemispheres were used for the comparison. (B) RT-qPCR analysis of the top downregulated genes in the brain extracts. n = 4 pairs of *Stag2^f/y^* and *Stag2^f/y^;Cre* littermates were used. Mean ± SD. (C) Heatmap of the expression of myelin-enriched genes that were down-regulated by more than two folds in *Stag2^f/y^;Cre* brains. *L1* and *R1*, left and right brain hemispheres of the *Stag2^f/y^#1* mouse. *L2* and *R2*, left and right brain hemispheres of the *Stag2^f/y^#2* mouse. *L1’* and *R1’*, left and right brain hemispheres of the *Stag2^f/y^;Cre #1* mouse. *L2’* and *R2’*, left and right brain hemispheres of the *Stag2^f/y^;Cre#2* mouse. The biological pathways of these genes are labeled on the right. (D) Top canonical pathways identified by ingenuity pathway analysis (IPA) of the DEGs.

Myelin is the membrane sheath that wraps around axons to facilitate rapid nerve conduction and maintain metabolic supply (Williamson and Lyons, 2018). Dynamic myelination in the central nervous system (CNS) is critical for proper neurodevelopment, and defective myelination is associated with autoimmune and neurodegenerative diseases (Mathys et al., 2019; Wolf et al., 2021). As cholesterol biosynthesis is essential for normal myelination (Hubler et al., 2018; Saher et al., 2005), we hypothesized that depletion of STAG2 caused myelination defects in the nervous system.

Indeed, brain sections of *Stag2^f/y^;Cre* mice showed greatly reduced luxol fast blue (LFB) staining compared to those of *Stag2^f/y^* and *Nestin-Cre* heterozygous mice (Figure 3A and S4A). Immunohistochemistry using antibodies against myelin proteins, MBP and PLP1, confirmed that *Stag2* cKO mice had significant defects in myelin fiber formation (Figure 3B-F). In both cerebral cortex and cerebellum, there were fewer and sparser myelin fibers in *Stag2^f/y^;Cre* mice, as compared to the *Stag2^f/y^* controls. Axon myelin ensheathment was further examined using transmission electron microscopy (Figure 3G). *Stag2^f/y^;Cre* mice had significantly fewer myelin-wrapped axons at optic nerves. Collectively, these data indicate insufficient myelination in the *Stag2* cKO mice. Myelination predominantly occurs at 3 weeks after birth in the mouse. The timing of premature death of *Stag2* cKO mice is thus consistent with defective myelination as a contributing factor to the lethality.

**Figure 3.**
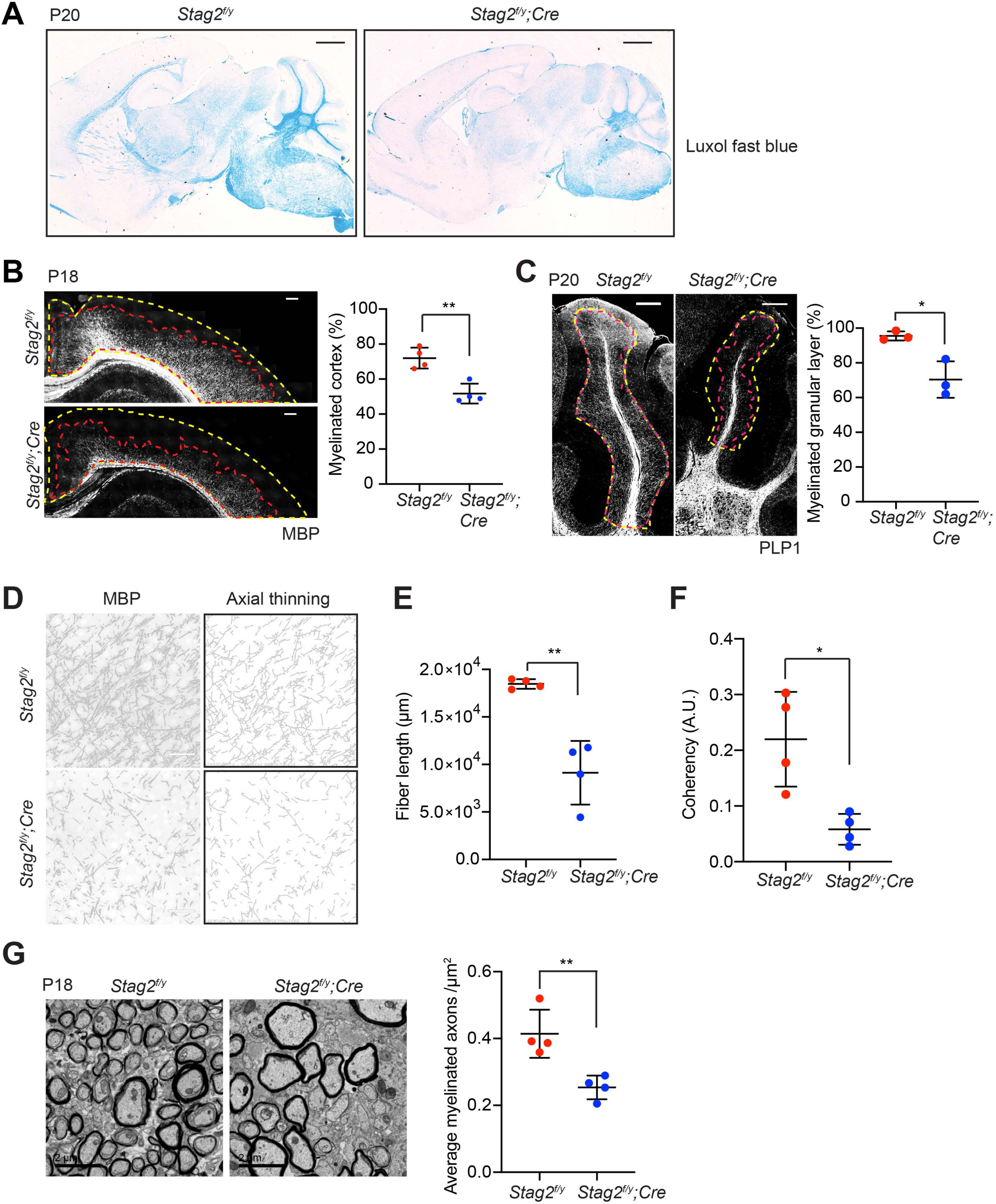
*Stag2* ablation in the nervous system compromises myelination during early postnatal development. (A) Luxol fast blue staining of the sagittal sections of *Stag2^f/y^* and *Stag2^f/y^;Cre* brains. n = 3 animals per genotype. Scale bar = 1 mm. (B) Immunohistochemistry staining with the anti-MBP antibody in the cerebral cortex (left panel). Antibody-stained areas and DAPI staining regions are marked with red and yellow dashed lines, respectively. Scale bar = 200 μm. Quantification of the percentage of the myelinated cortex is shown in the right panel. n = 4 pairs of *Stag2^f/y^* and *Stag2^f/y^;Cre* littermates were used (P18 or P21) for the comparison. **p < 0.01; Mean ± SD. (C) Immunohistochemistry staining with the anti-PLP1 antibody in the cerebellum (left panel). Antibody-stained areas and DAPI staining regions are marked with red and yellow dashed lines, respectively. Scale bar = 200 μm. Quantification of the percentage of the myelinated cerebellum granular layer is shown in the right panel. n = 3 pairs of *Stag2^f/y^* and *Stag2^f/y^;Cre* littermates were used (P20 or P25) for the comparison. *p < 0.05; Mean ± SD. (D) Higher magnification images (left panel) of the immunohistochemistry staining with the anti-MBP antibody in (B). Images processed through axial thinning are shown in the right panel. Scale bar = 50 μm. (E,F) Total fiber length (E) and fiber coherency (F) measured using the processed images in (D). n = 4 pairs of *Stag2^f/y^* and *Stag2^f/y^;Cre* littermates were used (P18 or P21). *p < 0.05, **p < 0.01; Mean ± SD. (G) Transmission electron microscopy images of the optic nerves (left panel). Scale bar = 2 μm. Quantification of myelinated axon distributions is shown in the right panel. n = 4 pairs of P18 *Stag2^f/y^* and *Stag2^f/y^;Cre* littermates were used. n ≥ 10 fields of each mouse were taken, and the average distribution of myelinated axons were calculated for each mouse and plotted. **p < 0.01; Mean ± SD.

### STAG2 regulates transcription in OLs

We examined *Stag1* and *Stag2* expression patterns in wild-type mouse brains by *in situ* hybridization using isotope-labeled RNA probes (Figure S4B). Both *Stag1* and *Stag2* were expressed at high levels in hippocampus, medial habenula, neocortex, and cerebellum granular layer. Aside from these regions, the *Stag2* transcripts were detected at relatively high levels in subventricular zone, thalamus, fiber tracts, midbrain, and hindbrain regions. *Stag2* is thus ubiquitously expressed in the brain.

Oligodendrocytes (OLs) are responsible for myelination in the CNS. To examine whether the OL lineage was affected by *Stag2* deletion, we performed single-cell RNA sequencing (scRNA-seq) analysis of *Stag2^f/y^;Cre* and *Stag2^f/y^* forebrains. As revealed by clusters in the t-SNE plot, the two genotype groups had similar cellular compositions (Figure 4A,B). All cell clusters were present in *Stag2^f/y^;Cre* brains, indicating generally normal neural cell differentiation. Cell-type identities were discovered with feature gene expression (Figure S5A). The OL lineage consisted of five clusters: cycling OL progenitors (OPCcycs), OL progenitors (OPCs), newly formed OLs (NFOLs), myelin-forming OLs (mFOLs), and fully matured OLs (MOLs). Quantification of the distributions of these five cell types within the OL lineage revealed a mild reduction in the number of MOLs in *Stag2^f/y^* forebrains (Figure 4C). We noticed that a higher percentage of neurons was recovered in the *Stag2^f/y^;Cre* group. Since the bulk RNA-seq results did not show global upregulation of neuron-specific genes, we suspect that neurons in *Stag2^f/y^;Cre* had fewer myelin-wrapped axons and were easier to be dissociated and kept alive during our library preparation for scRNA-seq. Thus, from the transcriptome analysis, we did not observe overt defects in most neural cell differentiation in the *Stag2*-deficient forebrain regions.

**Figure 4.**
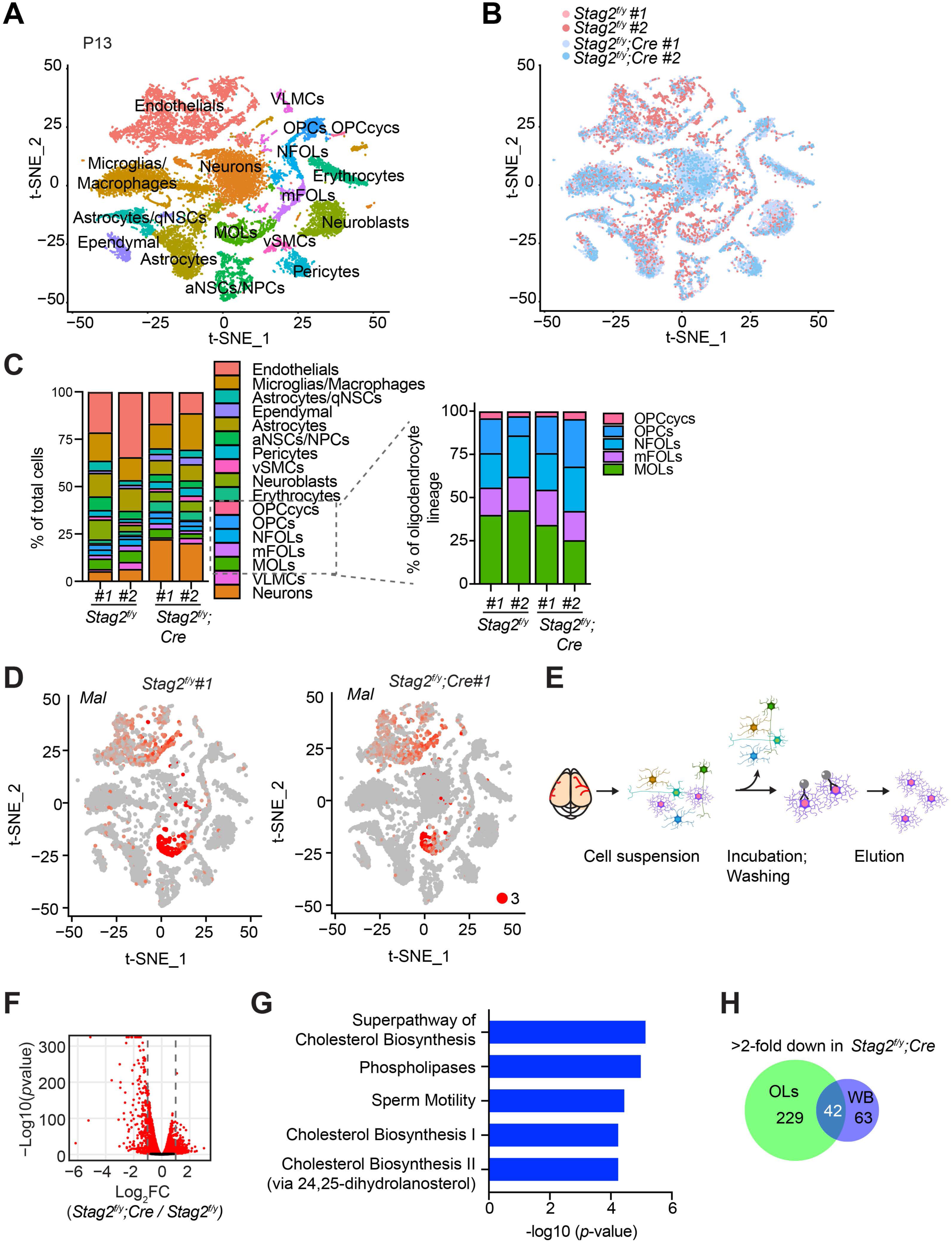
Deletion of *Stag2* in mouse brains causes differentiation delay and transcriptional changes in oligodendrocytes. (A) *t*-SNE plot of cell clusters in *Stag2^f/y^* and *Stag2^f/y^;Cre* forebrains analyzed by single-cell RNA-seq (scRNA-seq). n = 2 mice of each genotype were used in the scRNA-seq analysis. aNSCs/NPCs, active neural stem cells or neural progenitor cells; Astrocytes/qNSCs, astrocytes or quiescent neural stem cells; OPCcycs, cycling oligodendrocyte (OL) progenitor cells; OPCs, OL progenitor cells; NFOLs, newly formed OLs; mFOLs, myelin-forming OLs; MOLs, matured OLs; VLMCs, vascular and leptomeningeal cells; vSMCs, vascular smooth muscle cells. (B) *t*-SNE clustering as in (A) but colored by genotype. (C) Left panel: cell type composition and percentage as colored in (A). Right panel: percentage of cell clusters of the oligodendrocyte lineage. (D) FeaturePlot of a representative gene (*Mal*) specifically suppressed in MOLs of *Stag2^f/y^;Cre* forebrains. A maximum cutoff of 3 was used. (E) Experimental scheme of the magnetic-activated cell sorting (MACS) of primary OLs. (F) Volcano plot of bulk RNA-seq results of *Stag2^f/y^* and *Stag2^f/y^;Cre* primary OLs. (G) Top canonical pathways identified by IPA of the DEGs in (F). (H) Commons DEGs shared between bulk RNA-seq analyses of the whole brains (WB) and primary OLs.

We then performed trajectory inference and pseudotime analysis of the OL lineage (Figure S5B,C). Consistent with our cell-type assignment, pseudotime variables indicated continuous differentiation from OPCs to NFOLs, mFOLs, and MOLs (Figure S5D,E). The re-clustering of single cells in the OL lineage along the pseudotime path revealed that more cells were present in the terminal maturation stages in the *Stag2^f/y^* brains (Figure S5F,G). Conversely, more cells were retained at the undifferentiated stages in the *Stag2^f/y^;Cre* brains. Strikingly, some myelination genes, including *Mal*, were specifically repressed in *Stag2^f/y^;Cre* MOLs, with their expression in non-neural cells unaltered (Figures 4D and S6A). These observations suggest that STAG2 deficiency delays the maturation of OLs and compromises myelination-specific gene expression in mature OLs. Interestingly, compared to *Stag2* and genes encoding other cohesin core subunits, *Stag1* transcripts are less abundant in the OL lineage, except for cycling OPCs (Figure S6B,C). The low expression of *Stag1* in mature OLs might make these cells more dependent on *Stag2* for function.

To confirm the transcriptional defects in the OL lineage caused by *Stag2* deletion, we isolated primary OLs at intermediate differentiation stages from *Stag2^f/y^;Cre* and *Stag2^f/y^* forebrains with antibody-conjugated magnetic beads and conducted bulk RNA-seq analysis (Figure 4E). For both genotypes, the marker genes for NFOL and mFOLs were highly expressed in the isolated primary OLs (Figure S7A), suggesting that they mainly contained these two cell types. In *Stag2*-deleted OLs, 271 and 292 genes were downregulated or upregulated by more than two folds, respectively (Figure 4F). The top canonical pathway enriched in these DEGs was the cholesterol biosynthesis pathway (Figure 4G). Intriguingly, the downregulated genes were generally highly expressed in WT cells, whereas the upregulated genes had low expression levels in WT cells (Figure S7B-D). Among the 105 downregulated DEGs identified by RNA-seq analysis of the whole brain of *Stag2*-deficient mice, 42 were also differentially expressed in primary oligodendrocytes (Figure 4H). The cholesterol biosynthesis pathways were recognized as the major altered pathways (Figure S7E). Thus, defective cholesterol biosynthesis likely underlies hypomyelination and neurological defects in *Stag2* cKO mice.

We performed chromatin immunoprecipitation sequencing (ChIP-seq) experiments to examine the enrichment of the active transcription mark H3K27ac in *Stag2^f/y^* and *Stag2^f/y^;Cre* OLs and found that *Stag2* loss did not appreciably affect H3K27Ac enrichment at transcription start sites (TSS) (Figure S8A,B). Consistent with our RNA-seq results, the upregulated genes had much lower H3K27ac enrichment near their TSS, indicating that they were less active. We then checked the genomic distribution of STAG2 by ChIP-seq. Among other genomic loci, STAG2 was enriched at TSS of stable and downregulated genes, including genes in the cholesterol biosynthesis pathways (Figure S8C,D). It was enriched at the TSS of upregulated genes to a lesser extent, suggesting that *Stag2* loss might have indirectly affected the expression of these less active genes.

### *Stag2* deletion does not alter compartments or TADs in OLs

To investigate whether chromosome conformation was altered by *Stag2* deletion and whether that caused transcription dysregulation, we performed high-dimensional chromosome conformational capture (Hi-C) analysis of primary OLs isolated from *Stag2^f/y^* and *Stag2^f/y^;Cre* mice. We observed few compartment switching events in *Stag2*-deleted cells (Figure 5A-C and S9A-C). Virtually all genomic regions in *Stag2*-deleted cells were kept in their original compartment categories (AA or BB) (Figure 5C). Only a very small number of genomic regions switched compartments (AB or BA). Consistent with the RNA-seq data, analysis of average gene expression changes of DEGs in these genomic regions revealed that more genes located in the transcriptionally active A compartment (AA) were repressed in *Stag2*-deleted cells and more genes in the transcriptionally silent B compartment (BB) were upregulated (Figure 5D and S9D). Genes that switched from the A compartment to the B compartment were not more repressed compared to those that remained in the A compartment. Likewise, compared to genes that stayed in the B compartment, genes located in chromatin regions that switched from compartment B to A were not significantly activated. Acute depletion of all forms of cohesin eliminates TAD formation (Wutz et al., 2017). In contrast, deletion of *Stag2* had minimal impact on TAD formation in oligodendrocytes (Figure 5E-H, S9E,F), suggesting that STAG1-cohesin compensates for the loss of STAG2-cohesin in spatial organization of chromatin at larger than megabase scales. Therefore, our findings are inconsistent with compartment switching and TAD alterations being the underlying cause for the observed gene expression changes in STAG2-deficient OLs.

**Figure 5.**
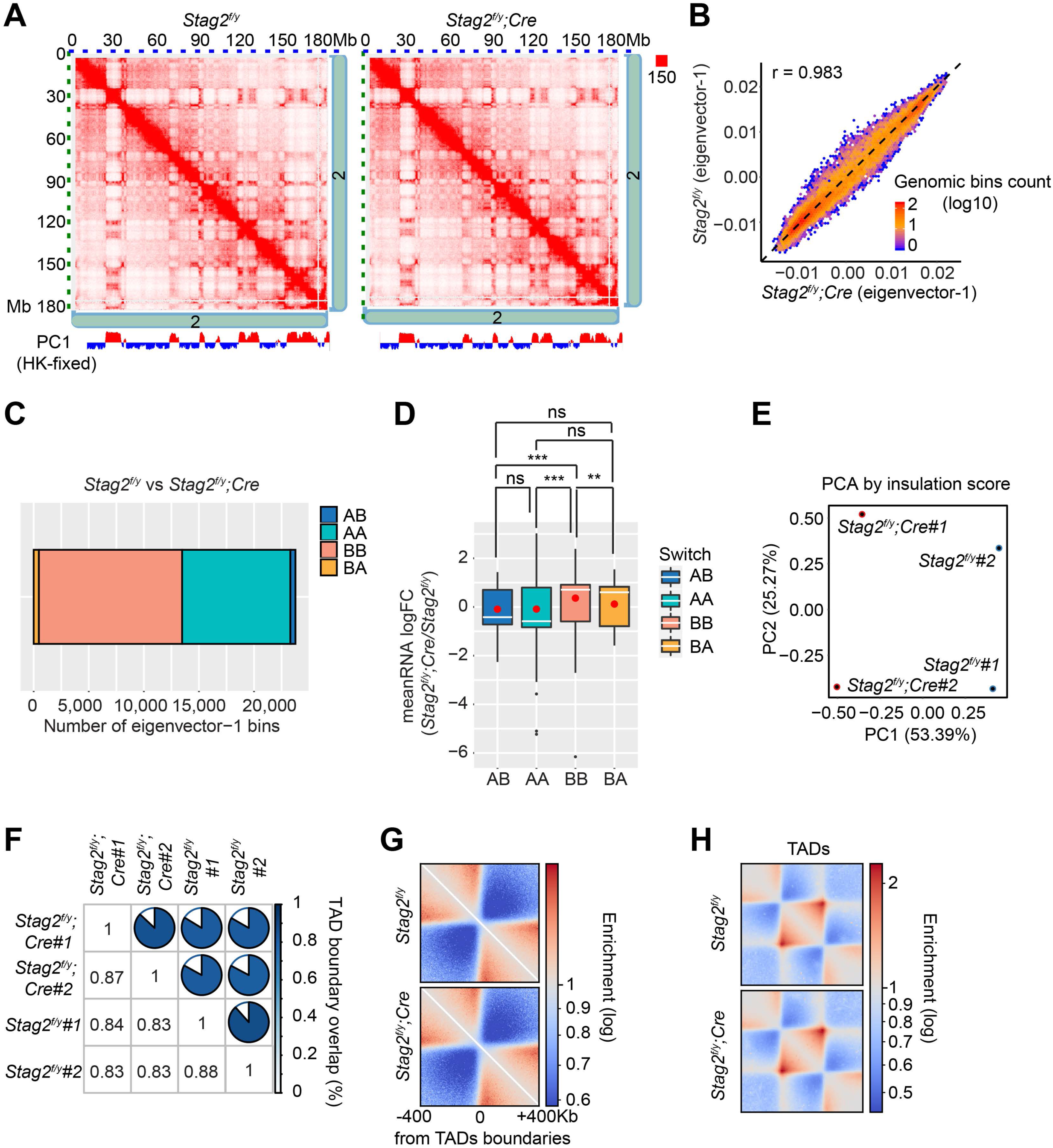
Loss of *Stag2* does not alter compartments and TADs in oligodendrocytes. (A) Representative snapshots of balanced Hi-C contact matrices of chromosome 2. Tracks of eigenvector-1 fixed with housekeeping genes are shown below, with A and B compartments shown in red and blue, respectively. (B) Hexbin plot of eigenvector-1 for genomic bins (100 kb) in *Stag2^f/y^* and *Stag2^f/y^;Cre* oligodendrocytes (OLs). (C) Chromatin bins were classified into four categories based on the eigenvector sign and whether it has switched with a delta bigger than 1.5. AB, changing from compartment A in *Stag2^f/y^* to compartment B in *Stag2^f/y^;Cre*; BA, from B in *Stag2^f/y^* to A in *Stag2^f/y^;Cre*; AA, A in both *Stag2^f/y^* and *Stag2^f/y^;Cre*; BB, B in both *Stag2^f/y^* and *Stag2^f/y^;Cre*. (D) Boxplot of averaged gene expression change of DEGs (RNA logFC cutoff of ± 0.58) inside each genomic bin. Bins counted: AA, 1646; AB, 56; BA, 69; BB, 910. Red dots represent the mean value. An unpaired Wilcoxon test was used for the statistical analysis. *p < 0.05; **p < 0.01; ***p < 0.001; ns, not significant. (E) Principal component analysis (PCA) plot of the insulation score. (F) Overlaps of TAD boundaries between genotypes and biological replicates. (G) Pile-up analysis of TAD boundary-centered local structure flanked by 400 kb chromatin regions. (H) Pile-up analysis of TAD local structure rescaled to an equal size.

### Promoter-anchored loops were reduced in *Stag2*-deleted OLs

While TAD boundaries are largely conserved among species and cell types, chromatin interactions within each TAD are more flexible and variable in cells undergoing differentiation, tumorigenesis, and reprogramming (Dixon et al., 2015; Dixon et al., 2012). Among the intra-TAD chromatin interactions, the enhancer-promoter loops are particularly important for transcription and are often cell-type specific. We examined whether chromatin loops in OLs were affected by *Stag2* loss. Compared to *Stag2^f/y^* OLs, *Stag2^f/y^;Cre* OLs had significantly fewer loops across almost all genomic distances (Figure 6A,B and S10). Loops specific to *Stag2^f/y^;Cre* OLs, which were likely mediated by STAG1-cohesin, were longer than STAG2-dependent *Stag2^f/y^*-specific loops. When genomic distances exceeded 0.25 Mb, the loops from *Stag2^f/y^;Cre* cells gradually gained higher scores over loops from *Stag2^f/y^* cells (Figure 6C). Therefore, STAG1-cohesin cannot completely compensate for STAG2-cohesin during loop formation. STAG1-cohesin-mediated loops are relatively longer than STAG2-cohesin-mediated loops, consistent with published findings in HeLa cells (Wutz et al., 2020).

**Figure 6.**
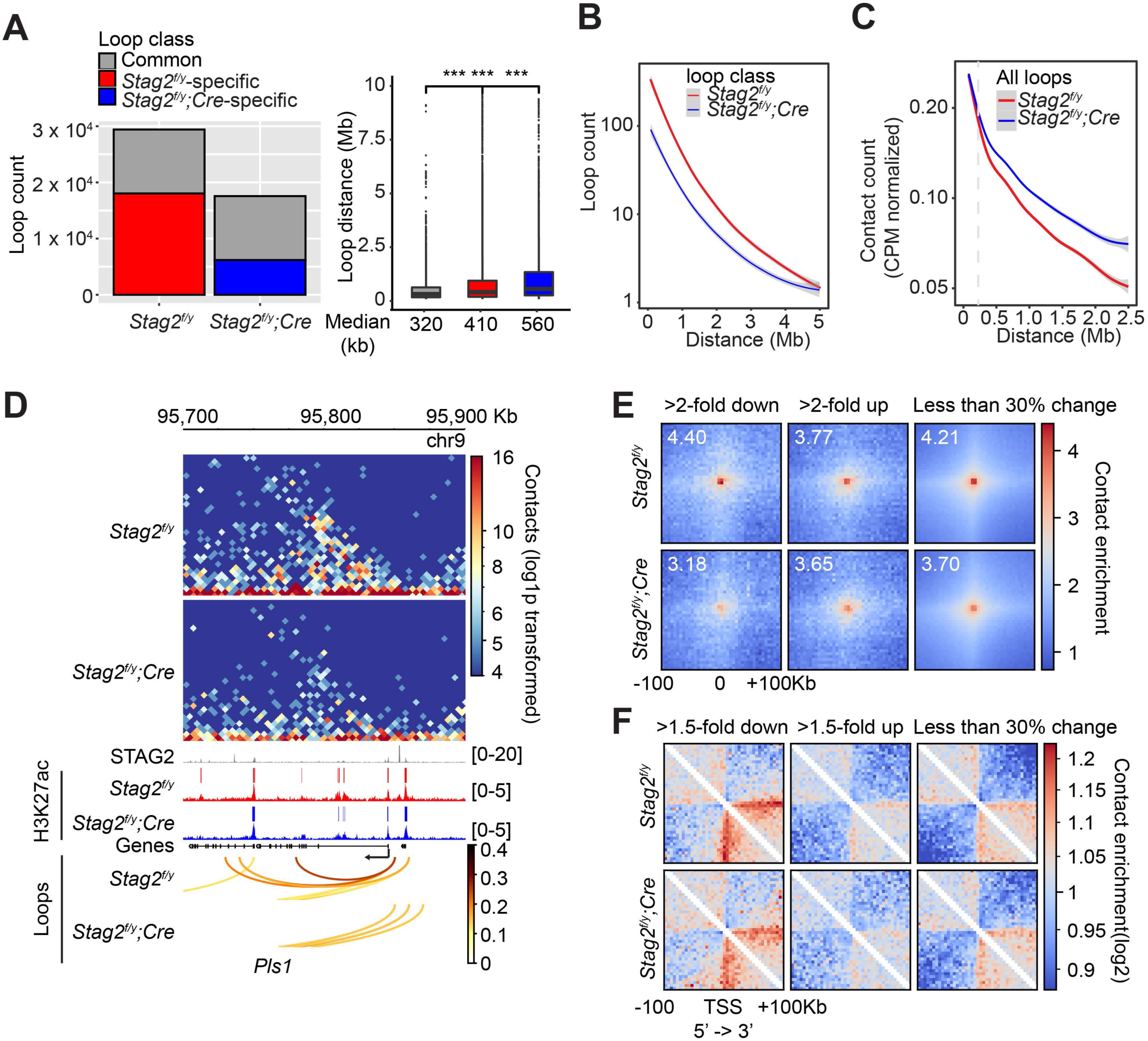
*Stag2* deletion impairs the formation of total and promoter-anchored loops in oligodendrocytes. (A) Loop counts (left panel) and length (right panel) in the indicated categories of *Stag2^f/y^* and *Stag2^f/y^;Cre* oligodendrocytes (OLs). (B) Loop counts plotted against loop length (from 0 to 5 Mb) of *Stag2^f/y^* and *Stag2^f/y^;Cre* OLs. (C) Normalized contact counts for loops across different genomic distances in *Stag2^f/y^* and *Stag2^f/y^;Cre* OLs. (D) Representative snapshots of contact maps at the *Pls1* gene locus. Tracks and narrow peaks from STAG2 and H3K27ac ChIP-seq as well as the loops are plotted below. Transcription direction is indicated by the black arrow. (E) Pile-up analysis of loop “dots”-centered local maps for the promoter-anchored loops of genes in the indicated categories. The maps are balanced, normalized by distance, and plotted at 5 kb resolution. The numbers indicate the enrichment of the central pixel over the upper left and bottom right corners. (F) Pile-up analysis of the local contact maps centered around the transcription start site (TSS) of genes in the indicated categories. Transcription directions are indicated below. 1,000 stable genes are chosen randomly and used for the analysis. The maps are balanced, normalized by distance, and plotted at 5kb resolution. Diagonal pixels are omitted.

We then tested whether the loop number decrease in *Stag2*-deficient cells could be a cause for transcriptional changes. When examining the local Hi-C maps, we noticed that loops anchored at gene promoters, including those of downregulated genes, were reduced in *Stag2^f/y^;Cre* oligodendrocytes (Figure 6D and S11). Promoter-anchored loops (P-loops) can potentially be promoter-promoter links, promoter-enhancer links, and gene loops. The total number of P-loops was proportionally decreased in *Stag2^f/y^;Cre* cells (Figure S12A). Moreover, the loops anchored at the downregulated genes were stronger than those at upregulated and stable genes (Figure S12B). We then compared P-loops associated with DEGs using pileup analysis of local contact maps. Loop enrichment at promoters of downregulated genes was reduced in *Stag2^f/y^;Cre* cells to a greater extent than that at promoters of upregulated and stable genes (Figure 6E). Taken together, our results suggest that *Stag2* loss diminishes short chromosome loops, including promoter-anchored loops. Highly expressed genes might be more reliant on these loops for transcription and are preferentially downregulated by *Stag2* loss.

We also performed pileup analysis of local chromatin regions flanking transcription start sites (TSS) (Figure 6F). Strikingly, we observed a clear stripe that extended from the TSS of downregulated gene only in the direction of transcription. The formation of promoter-anchored stripes (P-stripes) on aggregated plots is consistent with one-sided loop extrusion from the promoter to the gene body. The P-stripe was still present in *Stag2^f/y^;Cre* cells, suggesting that STAG1 could compensate for the loss of STAG2 and mediate its formation (Figure 6F and S12C).

## DISCUSSION

Cohesin is critical for the three-dimensional (3D) organization of the genome by extruding chromosome loops. Acute depletion of cohesin abolishes chromosome loops and TADs, but has moderate effects on transcription. The two forms of cohesin in vertebrate somatic cells, namely STAG1-cohesin and STAG2-cohesin, have largely redundant functions in supporting sister-chromatid cohesion and cell viability, but they have non-redundant functions during development. In this study, we have established a myelination-promoting function of STAG2 in the central nervous system (CNS) in the mouse. We further provide evidence linking hypomyelination caused by STAG2 loss to reduced promoter-anchored loops at myelination genes in oligodendrocytes.

### Myelination functions of STAG2 and implications for cohesinopathy

Selective ablation of *Stag2* in the nervous system in the mouse causes growth retardation, neurological defects, and premature death. STAG2 loss delays the maturation of oligodendrocytes and reduces the expression of highly active myelin and cholesterol biosynthesis genes in oligodendrocytes, resulting in hypomyelination in the CNS. Hypomyelination disorders in humans and mice are known to produce abnormal neurological behaviors similar to those seen in our *Stag2* cKO mice, suggesting that hypomyelination is a major underlying cause for the phenotypes in *Stag2* cKO mice. The growth retardation in these mice can be explained by insufficient secretion of growth hormones, which may be a consequence of defective neuronal signaling.

Mutations of cohesin subunits and regulators, including STAG2, cause the Cornelia de Lange syndrome (CdLS) and other similar developmental diseases, collectively termed cohesinopathy. CdLS patients exhibit short stature and developmental defects in multiple tissues and organs, including the brain. Although STAG2 mutations are implicated in human cohesinopathy, these mutations are rare and hypomorphic (Soardi et al., 2017). The cohesin loader NIPBL is the most frequently mutated cohesin regulator in cohesinopathy (Mannini et al., 2013). NIPBL deficiency is expected to affect the functions of both STAG1- and STAG2-cohesin. It is possible that the partial loss of STAG2-cohesin function leads to subtle myelination defects in patients with cohesinopathy. Indeed, lack of myelination in certain brain regions of CdLS patients has been reported (Avagliano et al., 2017; Vuilleumier et al., 2002). As myelination of the CNS mostly occurs after birth and during childhood, strategies aimed at enhancing myelination might help to alleviate certain disease phenotypes and symptoms.

### Mechanisms by which STAG2 promotes myelination

STAG2 promotes oligodendrocyte maturation and the expression of myelination genes in mature oligodendrocytes. Because STAG2 does not have an established cohesin-independent function, it most likely activates the myelination-promoting transcriptional program as a core component of cohesin. Consistent with previous reports (Rao et al., 2017), loss of STAG2-cohesin in oligodendrocytes does not affect genome compartmentalization, but reduces the number of relatively short chromosome loops, including promoter-anchored loops. Promoter-anchored loops at downregulated genes are reduced to a greater extent than those at stable and upregulated genes. These findings suggest that STAG2-cohesin promotes the myelination transcriptional program by forming promoter-anchored loops.

Pileup analysis of Hi-C maps reveals the formation of asymmetric promoter-anchored stripes in the direction of transcription at downregulated genes, indicative of active loading of cohesin at transcription start sites followed by one-sided loop extrusion from the promoter to the gene body. The stripes are, however, not reduced in STAG2-deficient cells. Because both forms of cohesin are capable of loop extrusion, it is possible that STAG1-cohesin can compensate for the loss of STAG2-cohesin in loop extrusion. It remains to be tested whether the intrinsic kinetics and processivity of loop extrusion mediated by the two forms of cohesin are differentially regulated by cellular factors or posttranslational modifications and whether these differences contribute to their non-redundant roles in transcription regulation.

We envision three possibilities that may account for why oligodendrocytes, but not other cell types, are more severely affected by *Stag2* loss in the CNS. First, STAG2-cohesin may be more abundant than STAG1-cohesin in post-mitotic OLs, making them more dependent on STAG2 for proper functions. Second, STAG1-cohesin preferentially localizes to CTCF-enriched TAD boundaries whereas STAG2-cohesin is more enriched at enhancers lacking CTCF (Kojic et al., 2018). Enhancers are critical for cell-type-specific gene transcriptional programs. To cooperate with the axonal growth during postnatal neurodevelopment, enhancer-enriched transcription factors induce timely and robust gene expression in oligodendrocytes for proper myelination (Mitew et al., 2014). The high-demand for enhancer function may render the transcription of myelination genes more reliant on STAG2-cohesin. Finally, the C-terminal regions of STAG1 and STAG2 are divergent in sequence and may bind to different interacting proteins and be subjected to differential regulation. STAG2 may interact with oligodendrocyte-specific transcription factors and be preferentially recruited to myelination genes. It will be interesting to investigate the interactomes of STAG1 and STAG2 in oligodendrocytes using mass spectrometry.

### STAG2-mediated chromosome looping and transcription

The mechanisms by which STAG2-dependent chromosome looping facilitates transcription are unclear at present. We propose several models that are not mutually exclusive (Figure 7). First, by forming promoter-enhancer loops, STAG2-cohesin brings the mediator complex and other enhancer-binding factors to the spatial proximity of the general transcriptional machinery at the promoter, thereby enhancing RNA polymerase II recruitment and transcription initiation. The existence of P stripes at STAG2-dependent genes in the Hi-C maps suggests that STAG2-mediated promoter-enhancer loops may involve enhancers located in the gene body. Second, loop extrusion by STAG2-cohesin may promote transcription elongation by regulating transcription-coupled pre-mRNA processing. For example, STAG2 has been shown to interact with RNA-DNA hybrid structures termed R-loops *in vitro* and in cells (Pan et al., 2020; Porter et al., 2021). R-loops formed between the nascent pre-mRNA and the DNA template impede transcription elongation and need to be suppressed (Moore and Proudfoot, 2009). When traveling with the transcription machinery on DNA, STAG2-cohesin might directly suppress R-loop formation or recruit other factors, such as the spliceosome, for co-transcriptional pre-mRNA processing and R-loop resolution. Third, STAG2-cohesin may establish promoter-terminator gene loops to recycle the RNA polymerase II that has finished one cycle of transcription back to the transcription start site for another round of transcription. Future experiments using high-resolution Hi-C methods in oligodendrocytes will allow us to better define the nature of STAG2-dependent promoter-anchored loops and stripes. It will also be interesting to examine whether *Stag2* deletion causes the accumulation of R-loops in downregulated genes and the incomplete splicing of their pre-mRNAs.

**Figure 7.**
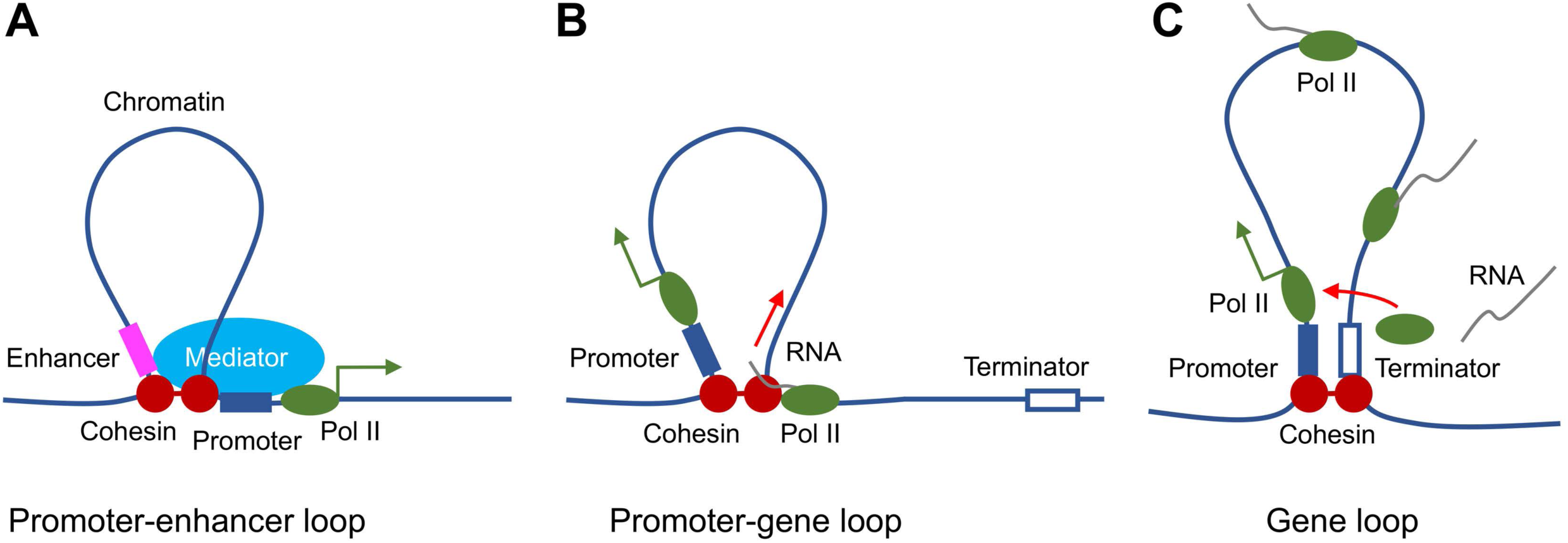
Proposed roles of STAG2-cohesin-mediated loop extrusion during transcription in oligodendrocytes. (A) STAG2-cohesin-mediated chromosome looping connects the enhancer and the promoter, thus facilitating interactions among oligodendrocyte-specific transcription factors, the mediator complex, and the general transcription machinery including RNA polymerase II. (B) STAG2-cohesin travels along the gene body via transcription-coupled loop extrusion to facilitate pre-mRNA processing. (C) STAG2-cohesin mediates the formation of gene loops that bring the terminator close to the promoter and facilitate Pol II recycling for multiple rounds of transcription.

## CONCLUSION

We have discovered a requirement for the cohesin subunit STAG2 in the myelination of the central nervous system in mammals. Our findings implicate hypomyelination as a contributing factor to certain phenotypes of cohesinopathy, including growth retardation and neurological disorders. We provide evidence to suggest that STAG2 promotes the myelination transcriptional program in oligodendrocytes through the formation of promoter-anchored loops. Our study establishes oligodendrocytes as a physiologically relevant cell system for dissecting the cellular functions and regulatory mechanisms of cohesin-mediated chromosome folding and genome organization.

## SUPPLEMENTAL INFORMATION

Supplemental Information including Supplemental Figures 1-12 can be found online.

## ACKNOWLEDGMENTS

We thank Sung Jun Bae for taking the mouse photos and John Shelton for help with histology and *in situ* hybridization. We are grateful to Jeffrey McDonald for the sterol composition analysis, Richard Lu and Lu Sun for providing reagents and advice for oligodendrocytes isolation, and Applied Bioinformatics Laboratories at NYU Langone Health for the Hi-C analysis. We also thank the Yu lab members for helpful discussions and for reading the manuscript critically. This study was supported by the Cancer Prevention and Research Institute of Texas (CPRIT) (RP160667-P2) and the Welch foundation (I-1441).

## AUTHOR CONTRIBUTIONS

N.C. and H.Y. conceived and designed the study. N.C. performed all the experiments. M.K., N.C., and C. X. performed the analysis and interpretation of NGS data. N.C. and B.M.E. performed histology and EM image analysis. N.C. and H.Y. wrote the manuscript with input from all authors. H.Y. supervised the project. All authors approved the manuscript.

## DECLARATION OF INTERESTS

The authors declare no competing interests.

## METHODS

### KEY RESOURCES TABLE

#### RESOURCE AVAILABILITY

##### Lead contact

Further information and requests for resources and reagents should be directed to and will be fulfilled by the Lead Contact, Hongtao Yu.

##### Material availability

Unique materials generated in this study will be available upon request.

##### Data and code availability

The RNA-seq, scRNA-seq, ChIP-seq, and Hi-C datasets generated and analyzed during the current study are available in the GEO repository, with the accession number xxx. Relevant code is available from the corresponding author upon request.

### EXPERIMENTAL MODEL AND SUBJECT DETAILS

#### Generation of *Stag2^f/f^* and *Stag2^f/y^* mice and mouse husbandry

The *STAG2* locus was targeted by inserting one neo cassette and two loxP sites flanking exon 8 via homologous recombination in the mouse embryonic stem (ES) cells. G418-selected positive ES clones were screened for successful targeting by nested PCR tests on both 5’ and 3’ integration sites of loxP. Four confirmed ES clones were then microinjected into mouse blastocysts. The chimeras were bred to the R26FLP mouse line for the removal of the neo cassette. *Stag2^f/+^* mice with the 129/B6 background were were crossed with *Stag2^f/y^* or wildtype C57BL/6J mice and maintained on this background. For the generation of the central nervous system-specific *Stag2^f/y^;Cre* mice, the *Stag2^f/f^* mice were crossed with the transgenic mice carrying one allele of *Cre* driven by the rat nestin promoter and enhancer (Tg(Nes-cre)1Kln, JAX stock #003771) (Giusti et al., 2014; Tronche et al., 1999).

All animals were handled in accordance with institutional guidelines of the Institutional Animal Care and Use Committee of University of Texas (UT) Southwestern Medical Center. All mice were housed in the antigen-free barrier facility with 12 hr light/dark cycles (6 AM on and 6 PM off). Mice were fed a standard rodent chow (2016 Teklad Global 16% protein rodent diet, Harlan Laboratories).

### METHOD DETAILS

#### Immunoblotting

The C-terminal fragment of human STAG2 protein was expressed and purified from *Escherichia coli* and used as the antigen to generate rabbit polyclonal antibodies against STAG2 at YenZym. Other antibodies were purchased from the following commercial sources: anti-SMC1 (Bethyl Laboratories, A300-055A), anti-SMC3 (Bethyl Laboratories, A300-060A), anti-RAD21 (Bethyl Laboratories, A300-080A), anti-SA1 (Bethyl Laboratories, A302-579A), anti-SA2 (Bethyl Laboratories, A302-581A), anti-α-TUBULIN (Sigma-Aldrich, DM1A), anti-MBP (Abcam, ab7349), anti-PLP1 (Abcam, ab28486), and anti-H3K27ac (Abcam, ab4729).

For immunoblotting, brain hemispheres were homogenized in a Precellys tissue homogenizer (Bertin Instruments) with the lysis buffer [20 mM Tris-HCl (pH 7.7), 137 mM NaCl, 2 mM EDTA, 10% (v/v) glycerol, 1% (v/v) TritonX-100, 0.5 mM dithiothreitol, 1 mM PMSF, 1 mM Na_3_VO4, 10 mM β-glycerophosphate, 5 mM NaF and protease inhibitors (Roche)]. Homogenized brain tissues were lysed on ice for 1 hr. The lysate was then subjected to centrifugation at 20,817 g at 4°C for 20 min and further cleared by filtering through a 0.45 μm filter. The cleared lysate was analyzed by SDS-PAGE and transferred to membranes, which was then incubated with the appropriate primary and secondary antibodies. The blots were imaged with the Odyssey Infrared Imaging System (LI-COR).

#### Tissue histology and immunohistochemistry

Mouse brains were fixed in 10% neutral buffered formalin solution for 48 hr followed by paraffin embedding and coronal or sagittal sectioning at 5 μm. Hematoxylin and eosin (H&E) staining and Luxol fast blue staining were performed by the Molecular Pathology Core at UT Southwestern Medical Center. Investigators were blinded to the genotype. Images were acquired with the DM2000 microscope (Leica) at 1.25X resolution.

Immunohistochemistry was performed as previously described (Choi et al., 2016). Briefly, deparaffinized sections were fixed with 4% paraformaldehyde, subjected to antigen retrieval by boiling with 10 mM sodium citrate (pH 6.0), and then incubated with the indicated antibodies at 1:100 dilution. The slides were scanned with an Axioscan.Z1 microscope (Zeiss) at 40X resolution at the Whole Brain Microscopy Facility at UT Southwestern Medical Center. Images were processed and quantified with Image J. For the myelinated fiber length measurement and coherency analysis, coronal sections of the brain cortex stained with the anti-MBP antibody were processed as previously described (van Tilborg et al., 2017). The myelinated axial thinning and fiber length measurement were performed by the plugin DiameterJ. The coherency analysis of myelinated axons was performed with the plugin OrientationJ.

#### Isolation of primary oligodendrocytes

The immunomagnetic isolation of oligodendrocytes from *Stag2^f/y^* and *Stag2^f/y^;Cre* P12-P14 pups was conducted using anti-O4 microbeads (Miltenyi Biotec) according to a published protocol (Flores-Obando et al., 2018). Briefly, dissected brain cortex was triturated in the papain dissociation solution with B-27 supplement. The single-cell suspension was incubated with anti-O4 magnetic beads and passed through a magnetic column to enrich oligodendrocytes. Freshly prepared oligodendrocytes were directly used or fixed for subsequent analysis.

#### Metabolic cage analysis

Mice were singly housed in shoebox-sized cages with a five-day acclimation period followed with a four-day recording period. Recorded parameters were analyzed by the TSE system and normalized to body weight. The experiments were conducted by the core personnel under the core protocol at the Metabolic Phenotyping Core at UT Southwestern Medical Center. Investigators were blinded to the genotype.

#### Growth hormone and IGF-1 detection

Blood samples were collected from facial bleeding without fasting. Plasma growth hormone levels were determined with the rat/mouse growth hormone ELISA kit (EMD Milipore, EZRMGH-45K). Plasma IGF-1 concentrations were measured using the mouse/rat IGF1 Quantikine ELISA kit (R&D Systems).

#### Sterol and oxysterol composition analysis

Brain hemispheres were pre-weighed and snap-frozen for extraction and measurement by mass spectrometry. The sterol extraction and quantitative analysis were conducted at the Center of Human Nutrition at UT Southwestern Medical Center as described previously (McDonald et al., 2012).

#### Electron microscopy

*Stag2^f/y^* and *Stag2^f/y^;Cre* P18 pups were transcardially perfused with 4% paraformaldehyde, 1% glutaraldehyde in 0.1 M sodium cacodylate buffer (pH 7.4). Tissues were dissected and fixed with 2.5% (v/v) glutaraldehyde in 0.1 M sodium cacodylate buffer (pH 7.4) for at least two hours. After three rinses with the 0.1 M sodium cacodylate buffer, optic nerve samples were embedded in 3% agarose and sliced into small blocks. All samples were again rinsed with the 0.1M sodium cacodylate buffer three times and post-fixed with 1% osmium tetroxide and 0.8 % potassium ferricyanide in the 0.1 M sodium cacodylate buffer for three hours at room temperature. Blocks were rinsed with water and *en bloc* stained with 4% uranyl acetate in 50% ethanol for two hours. Samples were dehydrated with increasing concentrations of ethanol, transitioned into propylene oxide, infiltrated with Embed-812 resin, and polymerized in a 60°C oven overnight. Blocks were sectioned with a diamond knife (Diatome) on a Leica Ultracut 7 ultramicrotome (Leica Microsystems) and collected onto copper grids, post-stained with 2% aqueous uranyl acetate and lead citrate. Images were acquired on a Tecnai G2 Spirit transmission electron microscope (Thermo Fischer) equipped with a LaB6 source using a voltage of 120 kV. Tissue processing, sectioning, and staining were completed by the Electron Microscopy Core at UT Southwestern Medical Center.

#### RNA-seq library preparation and sequencing

Total RNA was extracted from brain hemispheres or isolated oligodendrocytes with Trizol. RNA integrity was determined by the Agilent BioAnalyzer 2100. TruSeq Stranded mRNA library prep kit (Illumina) was used to generate the mRNA libraries. The libraries were analyzed by the Bioanalyzer and multiplexed and sequenced using the NextSeq 500 high output kit (400M reads) for the brain libraries or NextSeq 500 mid output kit (130M reads) for the isolated oligodendrocytes libraries at the Next Generation Sequencing Core at UT Southwestern Medical Center.

#### Differential expression and pathway analysis

Raw data from the sequencer were de-multiplexed and converted to fastq files using bcl2fastq (v2.17, Illumina). The fastq files were checked for quality using fastqc (v0.11.2) (Andrews, 2010) and fastq_screen (v0.4.4) (Wingett, 2011). Fastq files were mapped to the mm10 mouse reference genome (from iGenomes) using STAR (Dobin et al., 2013). Read counts were then generated using featureCounts (Liao et al., 2014). TMM normalization and differential expression analysis were performed using edgeR (Robinson et al., 2010). Pathway analysis was performed with the Ingenuity pathway analysis (IPA) software. Genes with more than 1.5-fold change and FDR < 0.01 were included in the brain RNA-seq pathway analysis. Genes with more than 2-fold change and FDR <0.05 were used for the pathway analysis of the RNA-seq data from oligodendrocytes.

#### RT-qPCR analysis

Single-stranded cDNAs were converted from 2 µg of total RNA extracted from mouse brains with the high-capacity cDNA reverse transcription kit (Applied Biosystems). Quantitative-PCR was conducted to determine transcript levels using gene-specific TaqMan probes (Applied Biosystems).

#### Single-cell RNA-seq

Single-cell suspension was prepared from forebrains of P13 *Stag2^f/y^* or *Stag2^f/y^;Cre* pups using the Papain Dissociation System (Worthington Biochemical, LK003150) according to the manufacturer’s instructions. Biological duplicates were made for each genotype. Single-cell RNA-seq libraries were generated with the Chromium Single Cell 3’ GEM, Library & Gel Bead Kit v3 (10x Genomics) according to the manufacturer’s guidelines. Cell density and viability were checked by the TC-20 Cell Counter (Bio-Rad). Cells were then loaded onto Chip B in the Chromium Controller (10x Genomics). 10,000 cells were targeted for each sample. The libraries were analyzed by the Bioanalyzer (Agilent) and pair-end sequenced in two flowcells of the NextSeq 500 High Output (400M) run. The sequencing was performed at the Next Generation Sequencing Core at UT Southwestern Medical Center.

Data de-multiplexing and alignment was performed using the Cell Ranger pipeline (https://support.10xgenomics.com/single-cell-gene-expression/software/pipelines/latest/using/ mkfastq) (10x Genomics). The raw features, barcodes, and matrixes were used as input for further analysis using the R package Seurat3 (Butler et al., 2018; Stuart et al., 2019) (https://satijalab.org/seurat/). Cells were filtered by the following criteria: nFeature_RNA (200-9500) and percent.mt < 10. After filtering, a total of 5,834 cells in *Stag2^f/y^#1*, 4,699 cells in *Stag2^f/y^#2*, 9,050 cells in *Stag2^f/y^;Cre#1*, and 3,073 cells in *Stag2^f/y^;Cre#2* were used for downstream analysis. 2,000 variable features were found from each normalized dataset. All datasets were then integrated using identified anchors (dims = 1:30). Standard scaling and principal component analysis (PCA), clustering (resolution = 0.5), and tSNE reduction (dims = 1:30) were performed on the integrated dataset. Cluster biomarkers were identified, and top features were examined. Clusters were then manually assigned to distinct cell type identities with knowledge from previous studies (Cahoy et al., 2008; Dulken et al., 2019; Marques et al., 2018; Marques et al., 2016; Marton et al., 2019; Saunders et al., 2018; Zeisel et al., 2018; Zywitza et al., 2018) (http://www.brainrnaseq.org/) (http://dropviz.org/). Clusters with the same cell type identities were merged. 5 clusters of oligodendrocyte lineage [cycling oligodendrocyte progenitors (OPCcycs), oligodendrocyte progenitors (OPCs), newly formed oligodendrocytes (NFOLs), myelin-forming oligodendrocytes (mFOLs) and fully matured oligodendrocytes (MFOLs)] were identified and selected for indicated gene expression comparison and plotting using Vlnplot or FeaturePlot functions. The trajectory analysis was performed using Monocle3 (Cao et al., 2019) in the oligodendrocyte cell population. Gene density plot over pseudotime was generated as previously described (Luecken and Theis, 2019).

#### ChIP-seq

Chromatin immunoprecipitation (ChIP) was performed as previously described (Liu et al., 2017). Briefly, isolated oligodendrocytes were fixed with 1% formaldehyde and fragmented with a sonicator (Branson 450). The fragmented chromatin was incubated with antibodies overnight at 4°C. Dynabeads Protein A (Thermo Fisher Scientific) was used for the immunoprecipitation. Libraries were generated by the Next Gen DNA Library Kit (Active Motif) with the Next Gen Indexing Kit (Active Motif) for STAG2 ChIP-seq or the KAPA HyperPrep Kits (KAPA Systems) for histone ChIP-seq. The libraries were analyzed by the Bioanalyzer and pool-sequenced with the NextSeq 500 mid output (130M) kit. After mapping reads to the mouse genome (mm10) by bowtie2 (v2.2.3) (Langmead and Salzberg, 2012) with the parameter “– sensitive”, we performed filtering by removing alignments with mapping quality less than 10 and then removing duplicate reads identified by Picard MarkDuplicates (v1.127). For STAG2 ChIP-seq, Picard MarkDuplicate was used to remove duplicates together with options to use molecular identifiers (MIDs) information in the reads. Enriched regions (peaks) were identified using MACS2 (v2.0.10) (Zhang et al., 2008), with a q-value cut-off of 0.05 for peaks. Peak regions were annotated by HOMER(Ross-Innes et al., 2012).

#### Hi-C library generation, sequencing, and analysis

Hi-C was performed at the Genome Technology Center at NYU Langone Health from 3.5-4.0 μg of DNA isolated from cells cross-linked with 2% formaldehyde at room temperature for 10 minutes. Experiments were performed in duplicates following the instructions from the Arima Hi-C kit (Arima Genomics, San Diego, CA). Subsequently, Illumina-compatible sequencing libraries were prepared by using a modified version of the KAPA HyperPrep library kit (KAPA BioSystems, Willmington, MA). Quality check steps were performed to assess the fraction of proximally ligated DNA labeled with biotin, and the optimal number of PCR reactions needed to make libraries. The libraries were loaded into an Illumina flowcell (Illumina, San Diego, CA) on a NovaSeq 6000 instrument for paired-end 50 reads.

Hi-C analysis was performed using the HiC-Bench pipeline (Lazaris et al., 2017; Tsirigos et al., 2012) (https://github.com/NYU-BFX/hic-bench). The read pairs were aligned and filtered with the following parameters: Genome-build=mm10; –very-sensitive-local –local; mapq=20; – min-dist 25000 –max-offset 500. The Juicer “pre” tool (Durand et al., 2016) (https:-//github.com/aidenlab/juicer) was used to generate the .hic file with default parameters. Sample duplicates were combined. The compartment analysis was done using the HOMER tool (Heinz et al., 2010) (http://homer.ucsd.edu/homer/index.html) with 100 kb bins. H3K27ac ChIP-seq data was used to assign A/B compartments. Eigenvector-1 bins were considered shifted (AB and BA) when the bin sign changed and the delta value was greater than 1.5. Topologically associated domains (TAD) and boundaries were identified with the HiCRatio method with the follow parameters: –min-lambda=0.0 –max-lambda=1.0 –n-lambda=6 –gamma=0 –distance=500kb – fdr=0.1. The .hic files were converted to .cool format for visualization and plotting with pyGenomeTracks (Lopez-Delisle et al., 2020) at 5 kb resolution.

#### Loop analysis and RNA-seq integration

The loops were classified into group-specific loops and common loops by using the significance cutoffs provided by Fit-HiC (Ay et al., 2014). A qvalue cutoff of 0.01 was used to identify significant loops in both groups. A loop is considered “group-specific” if it is only present in one group with a qval < 0.01 and not present in the other group with cutoff of qval < 0.1. Loop anchors were annotated with the gene promoter information (promoter defined as +/– 2 kb from the transcription start site). The genes were classified into “down” and “up” regulated genes using an FDR cutoff of 0.05, logFC cutoff of +/-0.58 and logCPM > 0. “stable” or less changed genes are defined as logFC < 0.38, and logCPM > 0. Random 1,000 genes were chosen for analysis and plotting. The genes were also grouped in “high”, “mid” and “low” expression groups by separating the genes in three quantiles according to the logCPM values. For the loop enrichment scores, normalized contact scores were computed using Fit-HiC at 10 kb resolution and bias corrected. Pileup analysis was performed with coolpup.py (Flyamer et al., 2020) with the KR method to balance the weight and random shift controls for distance normalization at 5 kb.

### QUANTIFICATION AND STATISTICAL ANALYSIS

Statistical analysis was conducted in GraphPad Prism or R-studio. Specific statistical methods and approaches are indicated for each figure in the figure legend.

## KEY RESOURCES TABLE

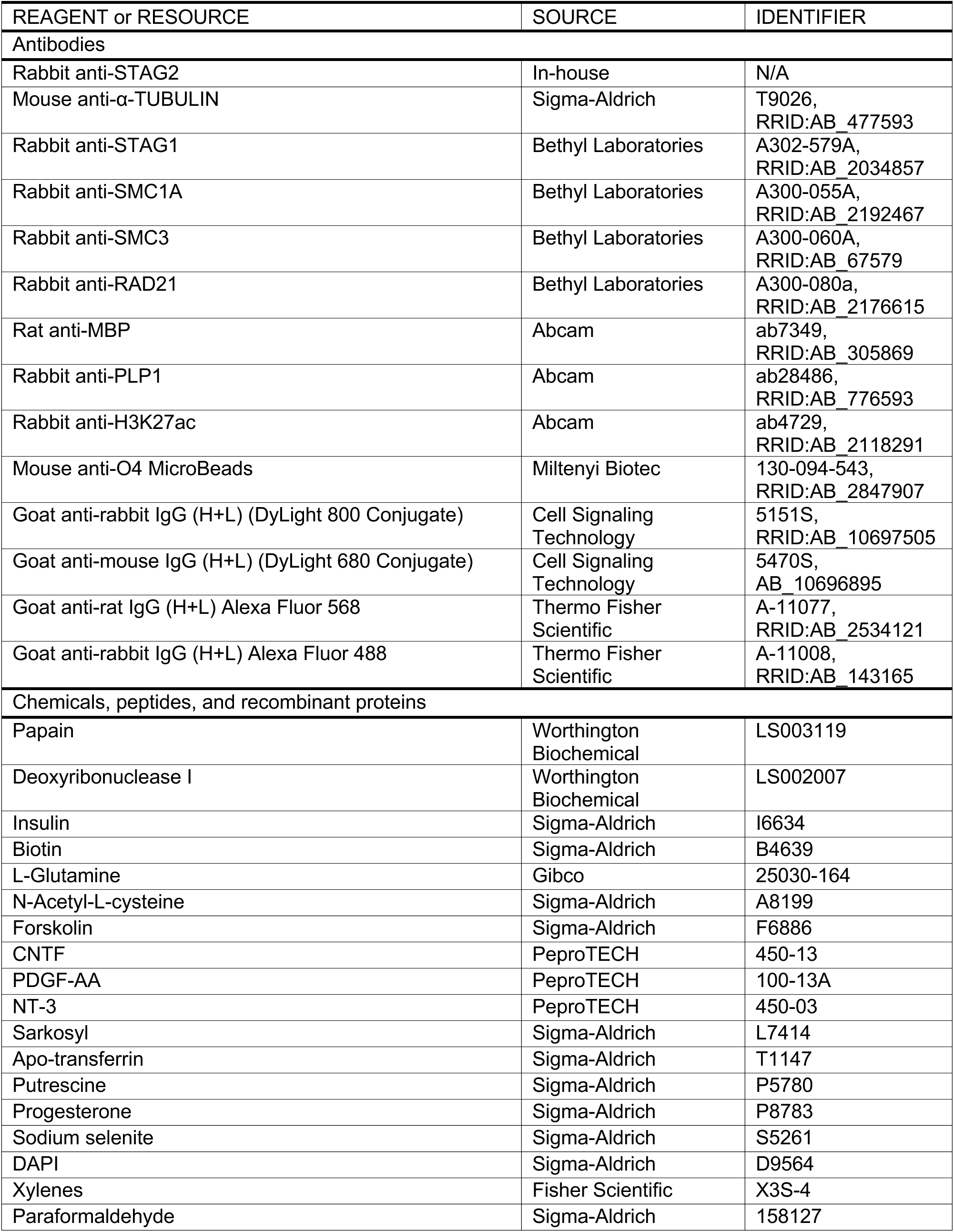
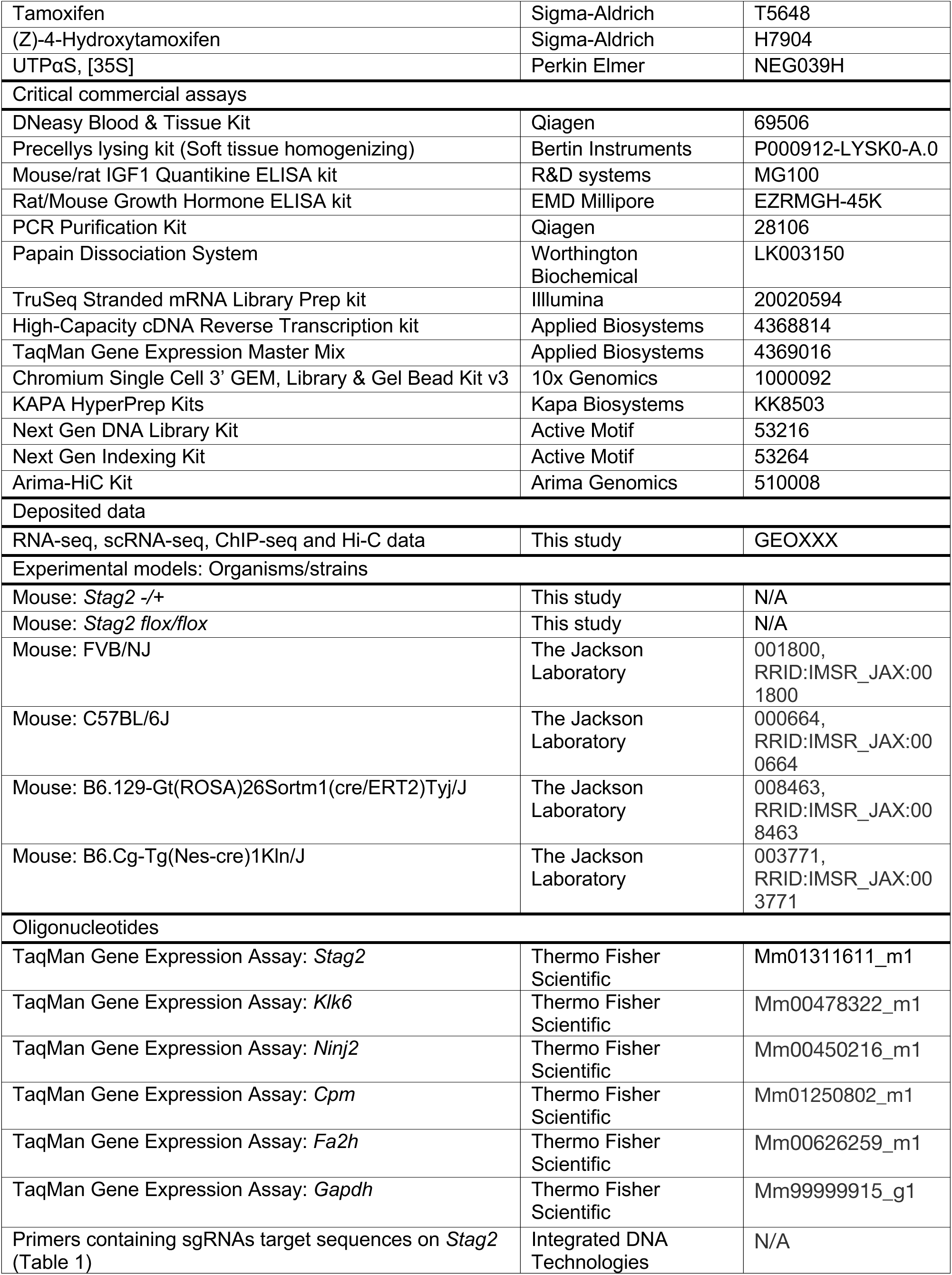
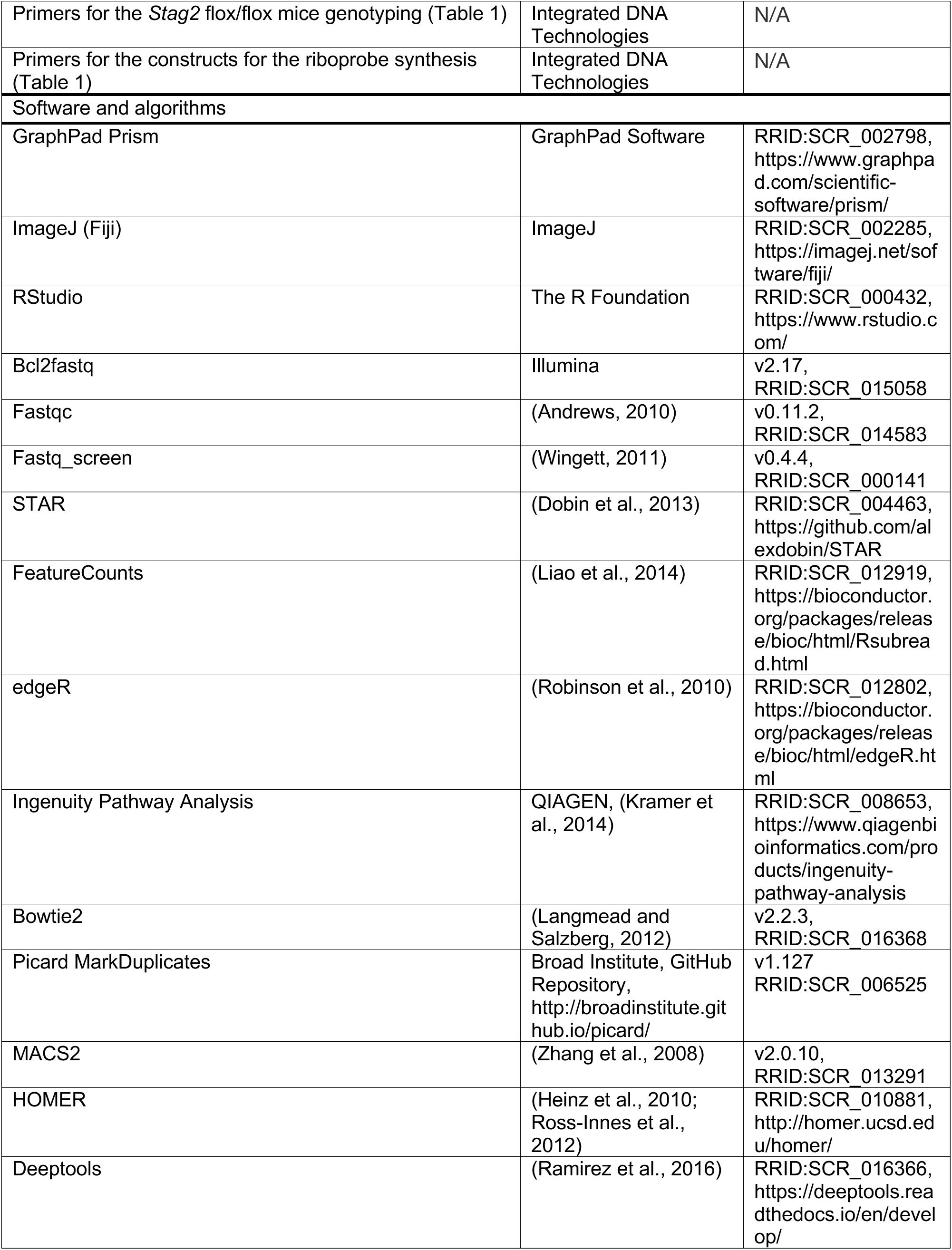
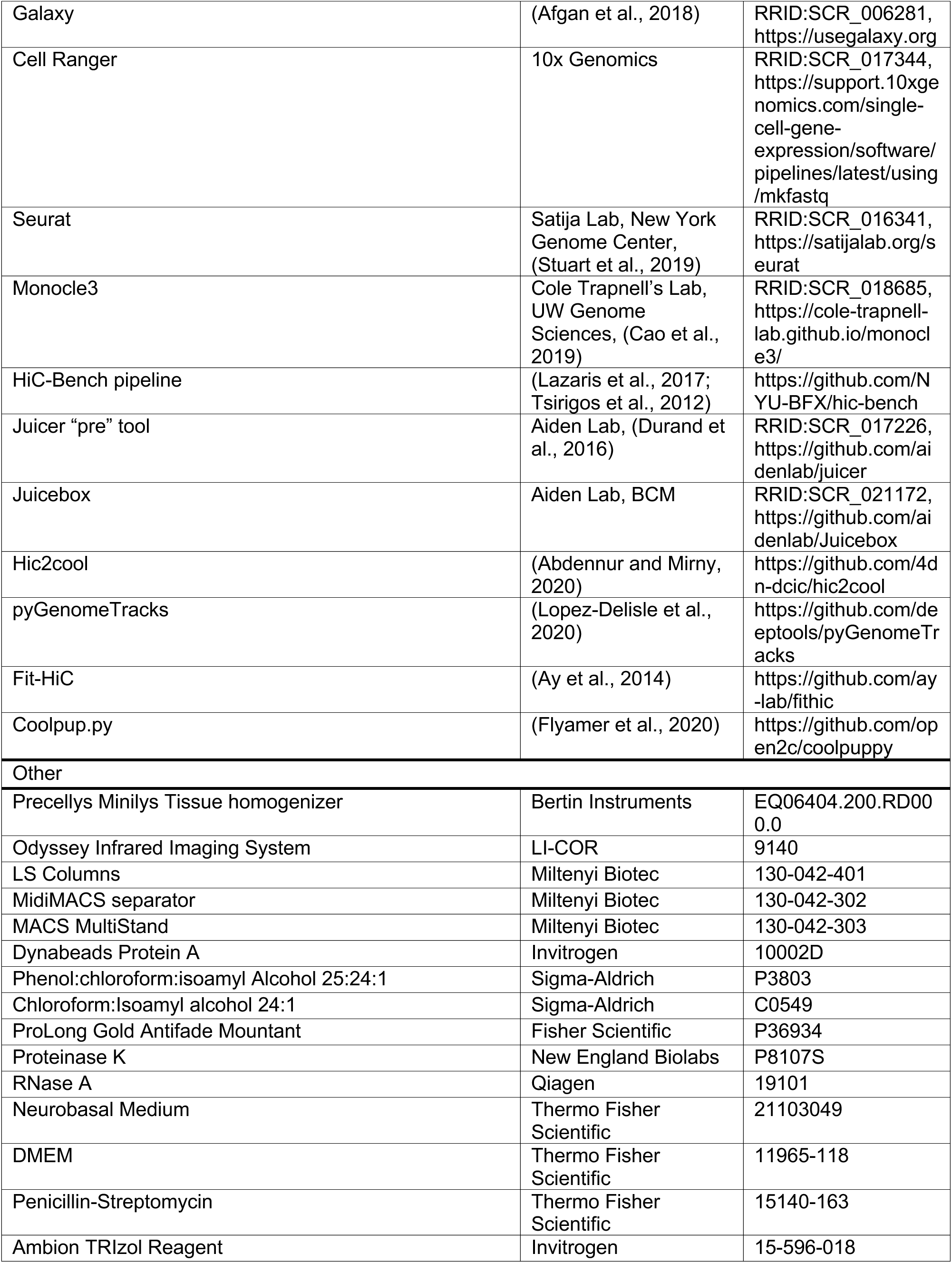
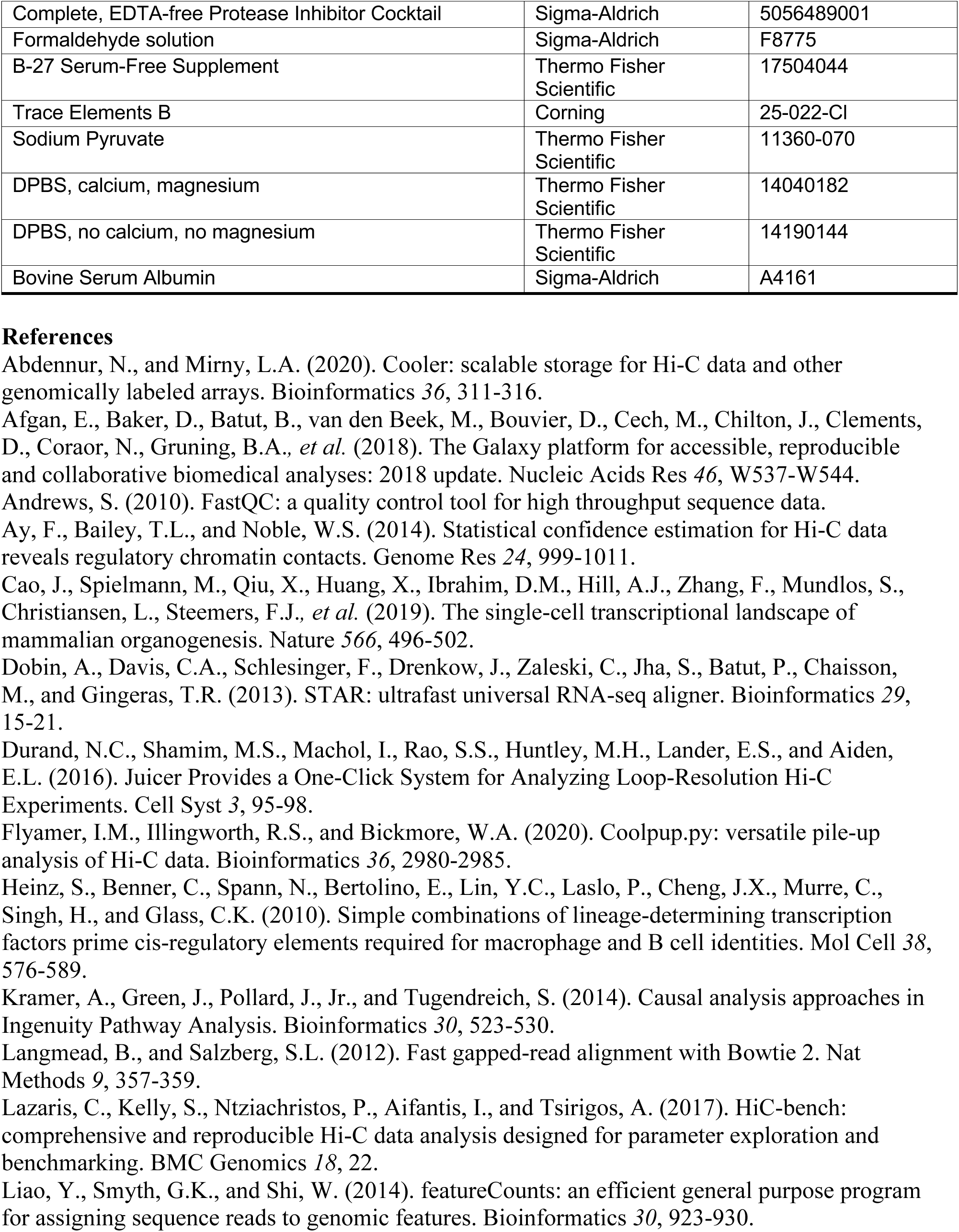
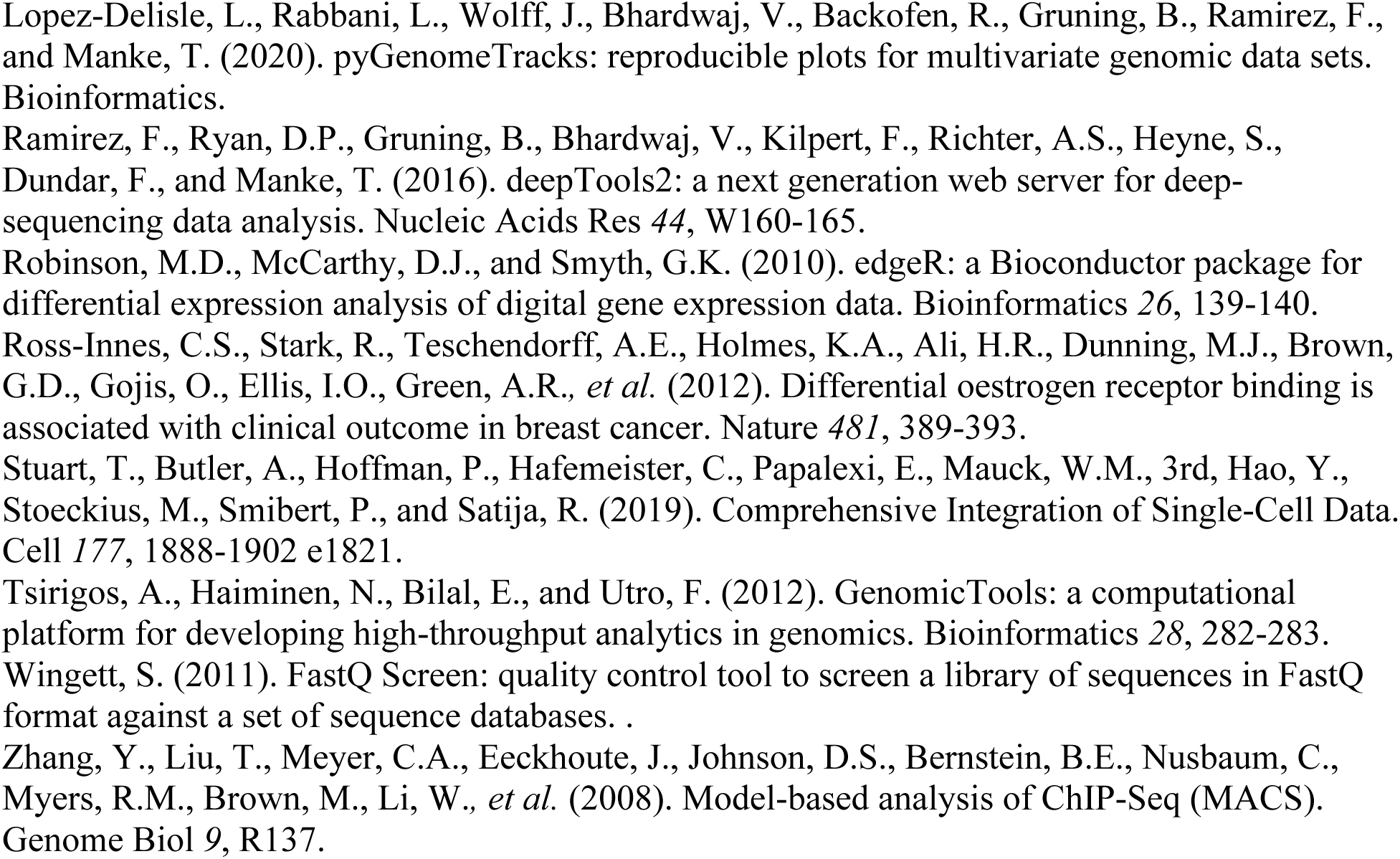

## Supplemental Information

**Figure S1.**
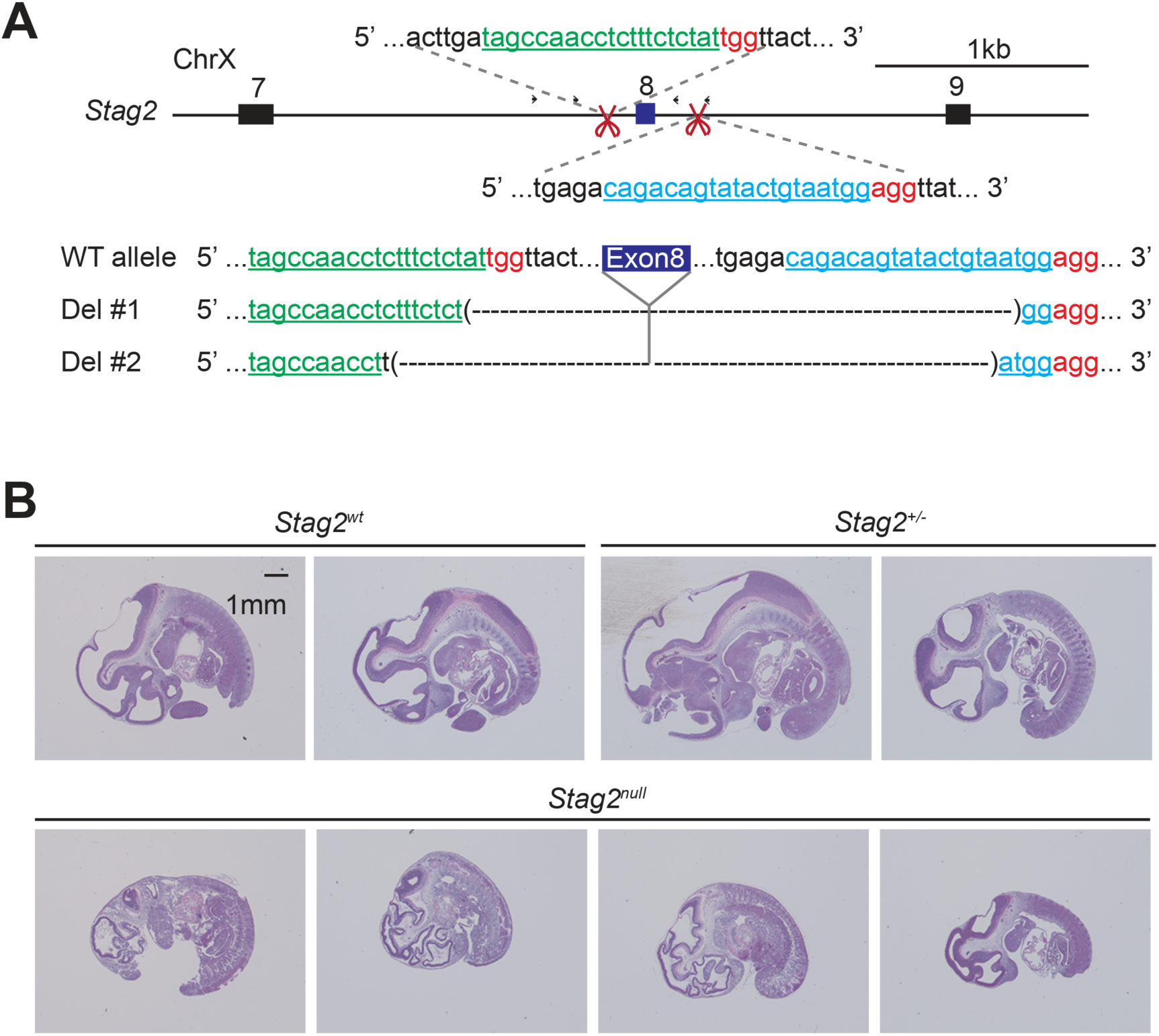
Generation of *Stag2* knockout mice using the CRISPR/Cas9 method. (A) Scheme for disrupting *Stag2* in the mouse genome using CRISPR/Cas9 with guide RNAs flanking exon 8. Sequencing analysis of the genomic DNA extracted from two *Stag2*-disrupted founder mice is shown below. (B) Hematoxylin and eosin (H&E) staining of sagittal sections of F2 embryos derived from F1 in (A) at E11.5.

**Figure S2.**
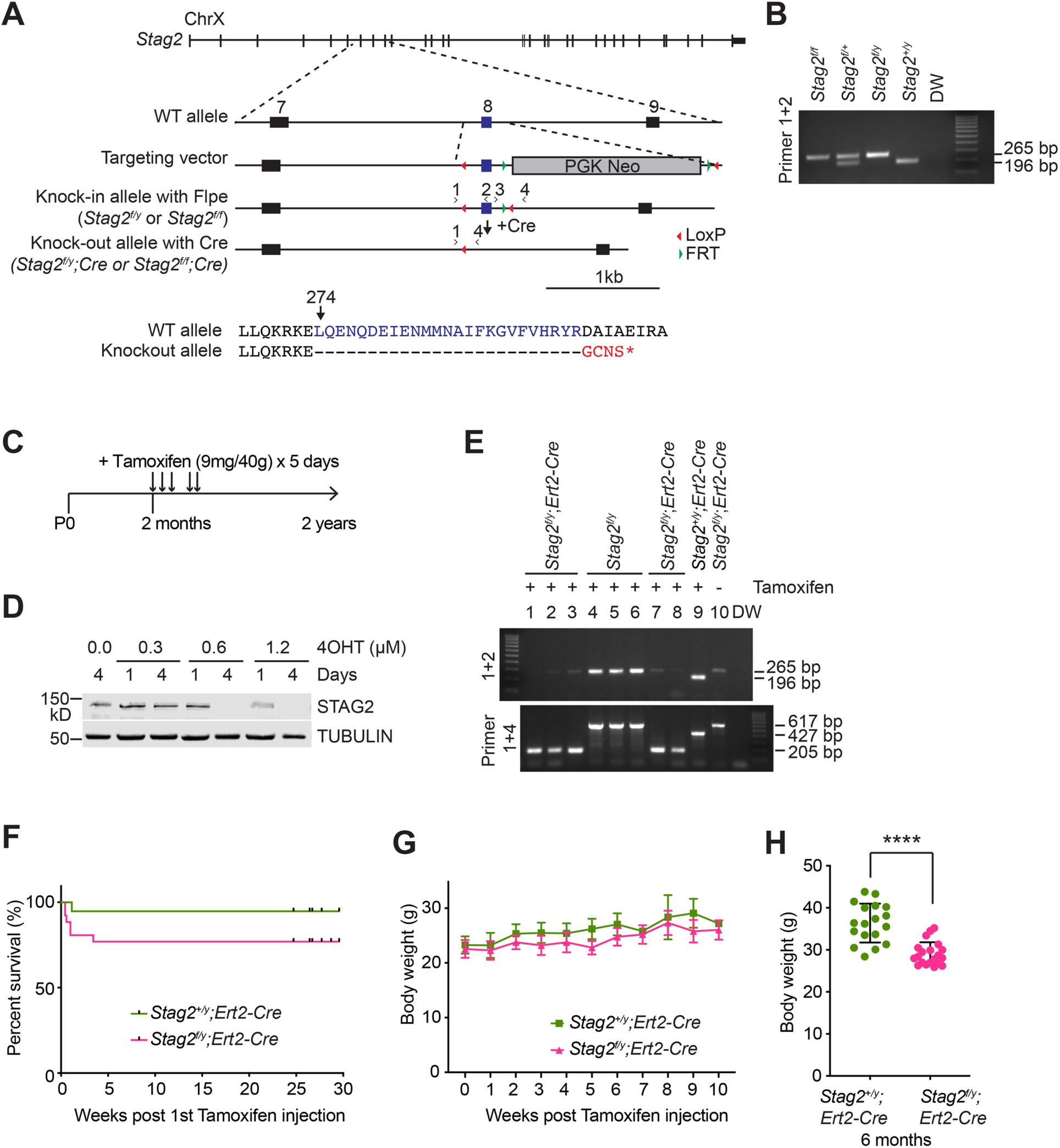
Generation of *Stag2* conditional knockout mice by gene targeting. (A) Scheme for creating the “floxed” *Stag2* allele by gene targeting. The genomic structure of the wild-type (WT) *Stag2* locus, the targeting vector, the knock-in allele, the disrupted allele after Cre-mediated recombination, and the positions of the genotyping primers are shown. The amino acid sequence of the knockout allele in the targeted region is shown and aligned with that of the WT allele. (B) PCR analysis of the genomic DNA extracted from the tails of indicated mice with the primers in (A). (C) Experimental scheme of tamoxifen injection into adult *Stag2^f/y^;Ert2-Cre* and *Stag^+/y^;Ert2-Cre* mice. (D) Western blotting of cell extracts from *Stag2^f/y^;Ert2-Cre* mouse embryonic fibroblasts (MEF) treated with or without 4-hydroxytamoxifen (4OHT). (E) PCR analysis of the genomic DNA extracted from the blood of indicated mice with the primers in (A). (F) Survival curves of *Stag2^f/y^;Ert2-Cre* (n=26) and *Stag^+/y^;Ert2-Cre* (n=19) mice after tamoxifen injection. (G) Body weight of mice in (F). (H) Body weight of *Stag2^f/y^;Ert2-Cre* (n=20) and *Stag^+/y^;Ert2-Cre* (n=18) mice at 6 months post tamoxifen injection. ****-p<0.0001.

**Figure S3.**
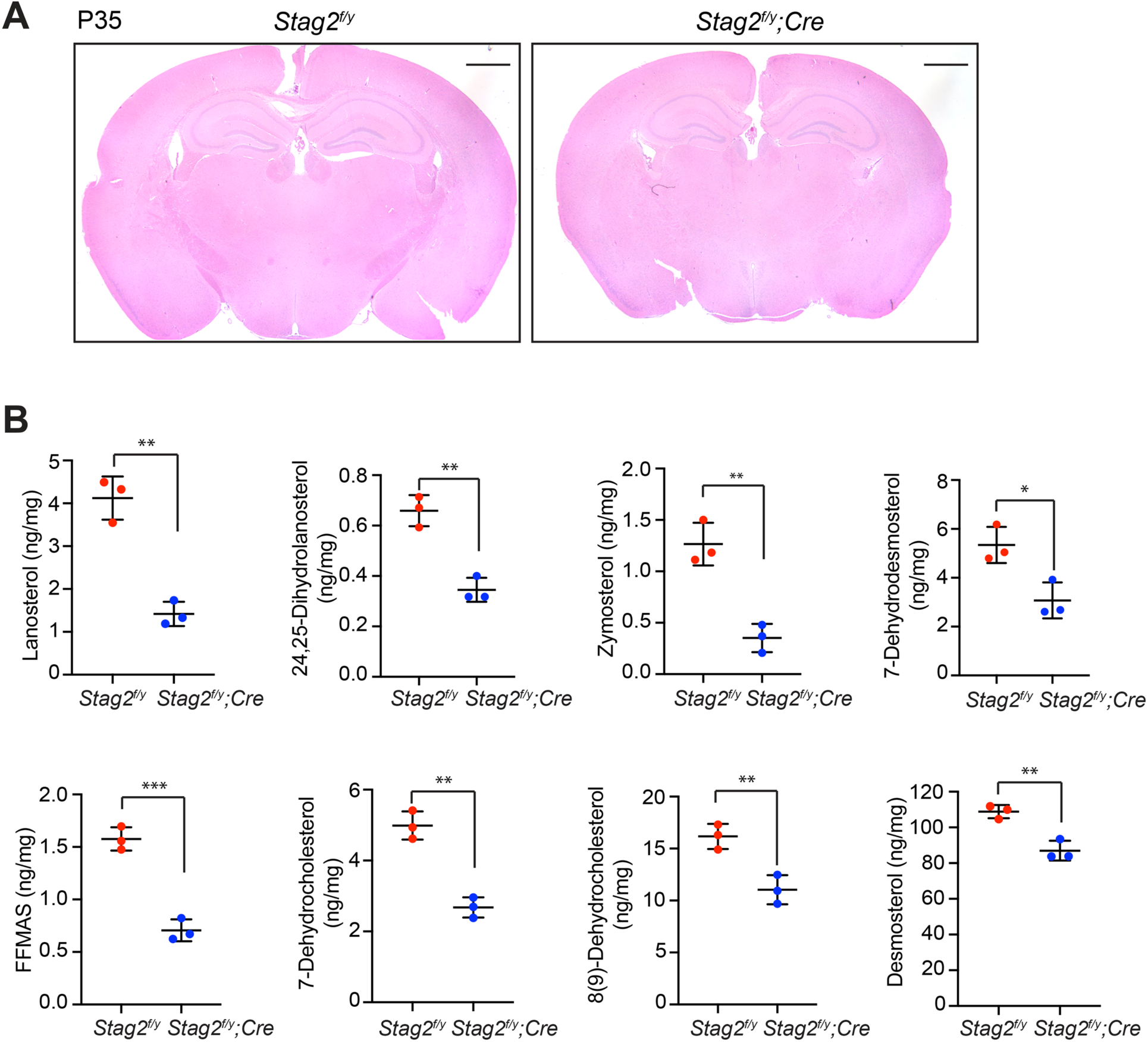
STAG2 deficiency in mouse brains attenuates cholesterol biosynthesis. (A) H&E staining of the coronal sections of *Stag2^f/y^* and *Stag2^f/y^;Cre* mouse brains. Scale bar, 1 mm. (B) Mass spectrometry analysis of cholesterol precursors in *Stag2^f/y^* and *Stag2^f/y^;Cre* brains. n = 3 mice per genotype. *p < 0.05, **p < 0.01, ***p < 0.001; Mean ± SD.

**Figure S4.**
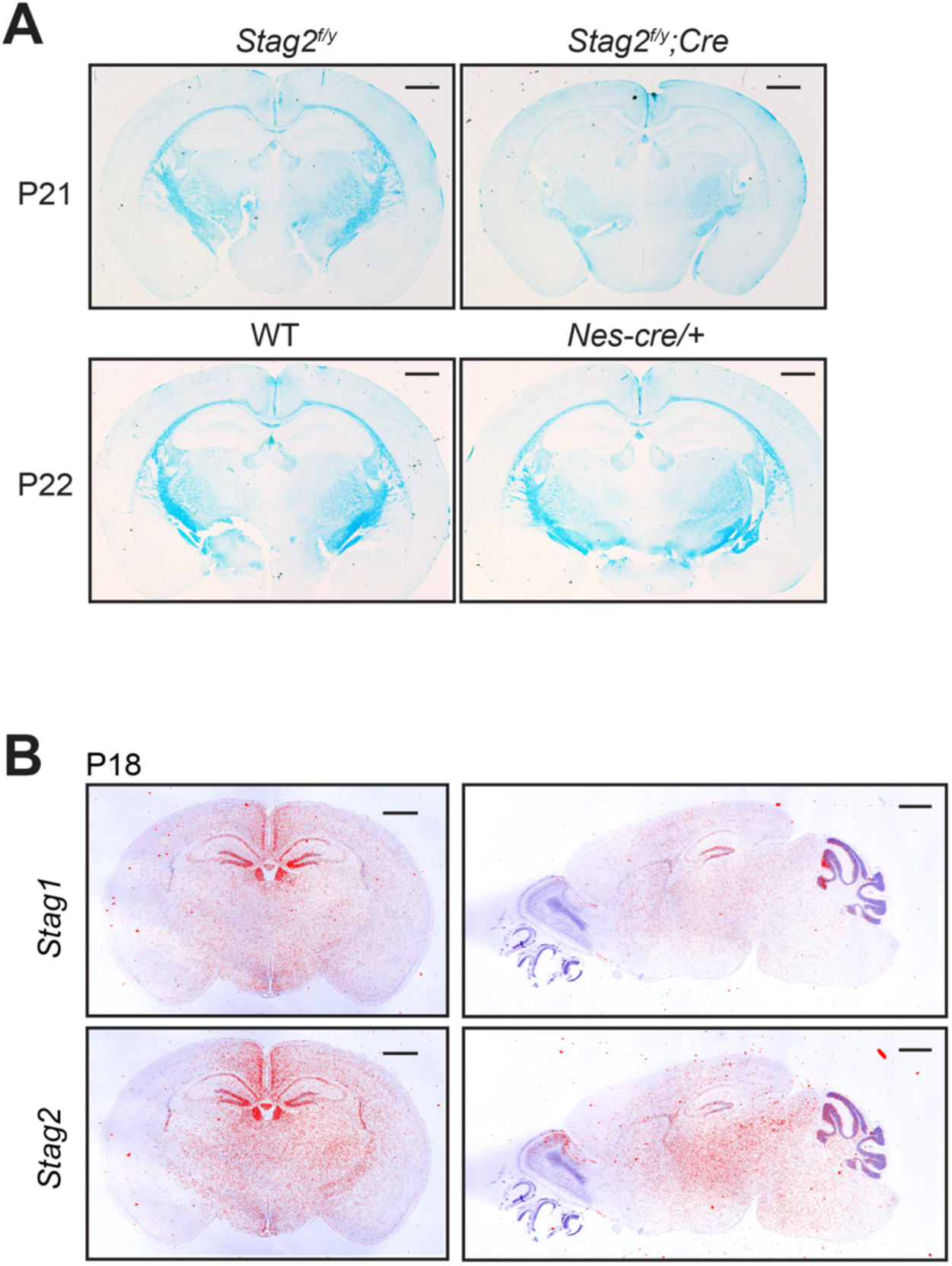
Brain-specific *Stag2* deletion impairs CNS myelination. (A) Luxol fast blue staining of the coronal brain sections of mice with the indicated genotypes. n = 3 mice each for *Stag2^f/y^* and *Stag2^f/y^;Cre* genotypes. n = 2 mice each for WT and *Nes-Cre/+* groups. animals per genotype. Scale bar = 1 mm. (B) In situ hybridization of ^35^S-labeled RNA probes of the coronal (left) and sagittal (right) sections of WT mouse brains. Bright field images (purple) were overlaid with autoradiography images (red). Scale bar = 1 mm.

**Figure S5.**
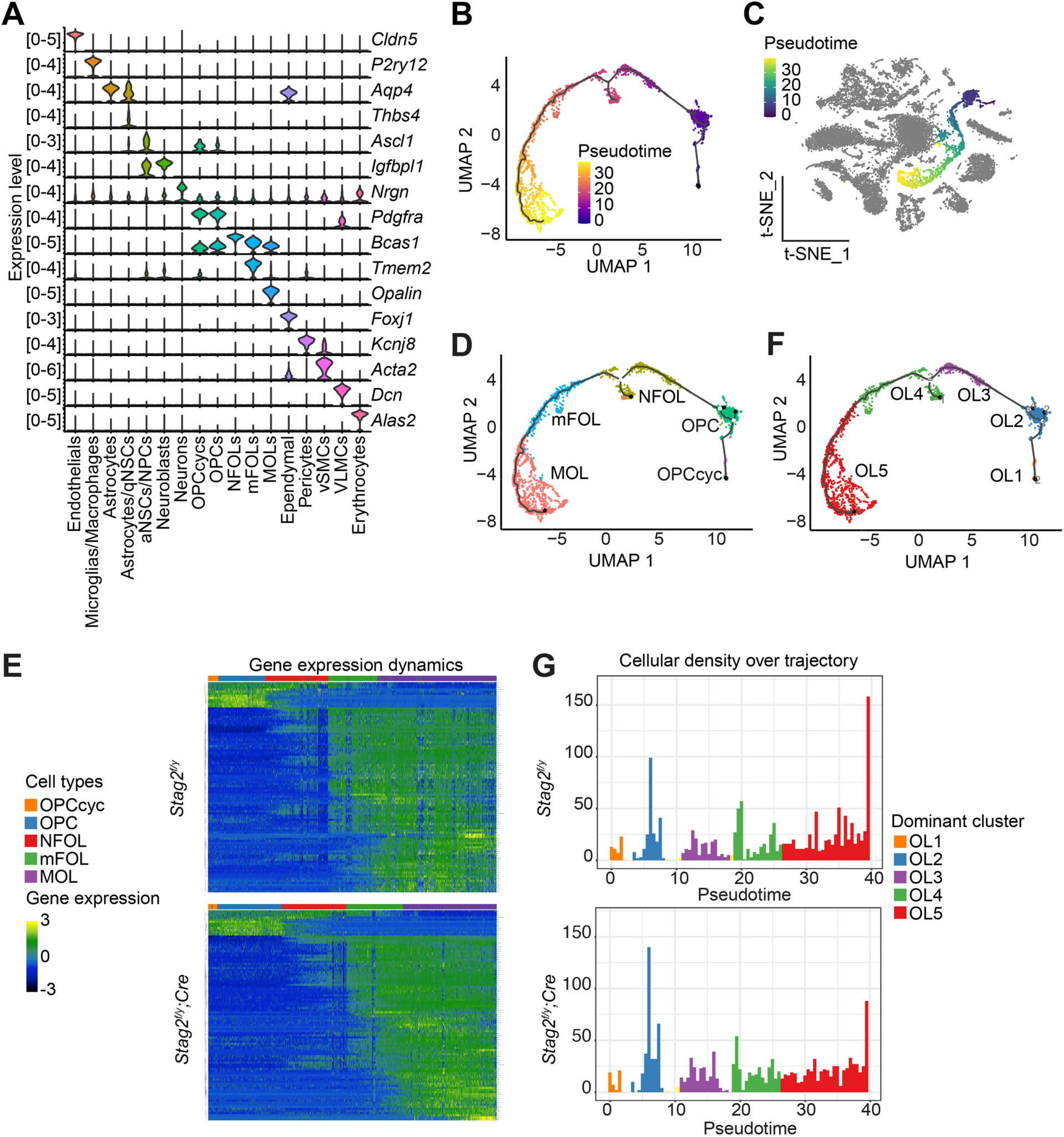
*Stag2* deletion causes differentiation delay in the oligodendrocyte lineage. (A) Violin plot of the expression levels of feature genes of the indicated brain cell types. (B) Trajectory inference analysis of oligodendrocyte (OL) lineage cells extracted from the single-cell RNA-seq dataset using Monocle3. Cells are colored from purple to yellow by pseudotime variables. (C) OL differentiation trajectory in the *t-SNE* plot. The OL lineage is colored from navy blue to yellow by pseudotime variables. Cells of other lineages are colored grey. (D) Distribution of the assigned OL cell types along the trajectory. (E) Heatmap of gene expression dynamics over pseudotime along the OL differentiation trajectory. Each row represents one of the top 100 most variable genes along pseudotime. Each column represents a single cell. (F) Re-clustered OL subgroups in the trajectory inference analysis. (G) Cell density across pseudotime for the OL differentiation trajectory. Dominant clusters for each pseudotime bin are color labeled as in (F).

**Figure S6.**
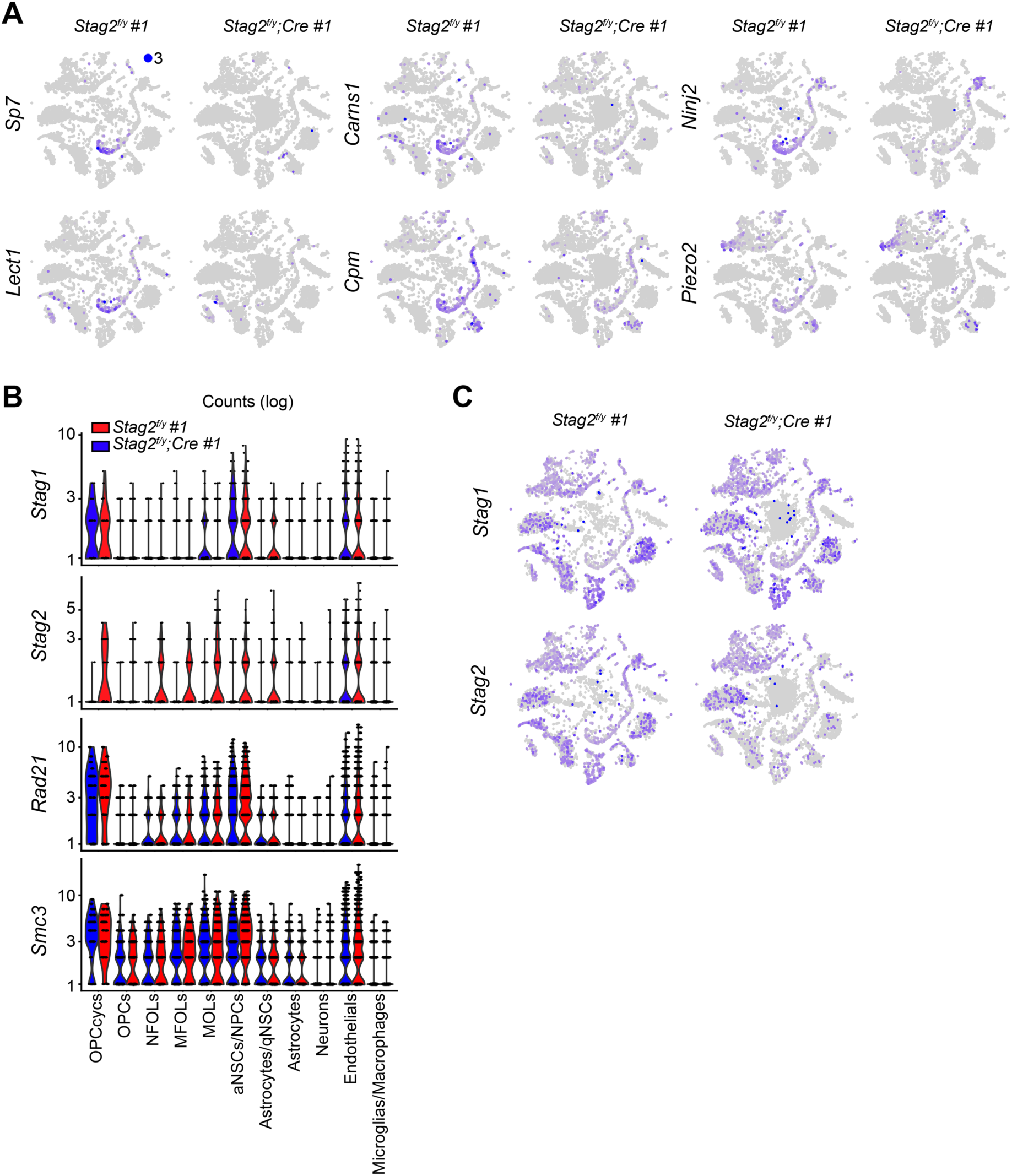
STAG2 regulates the transcription of oligodendrocyte genes. (A) FeaturePlot of the expression levels of representative down-regulated genes in the *Stag2^f/y^;Cre* whole brains. Maximum cutoff of 3 was used. (B) Violin plot of the expression of cohesin subunit genes in the indicated brain cell types from the scRNA-seq transcriptome analysis. (C) FeaturePlot of the expression of *Stag1* and *Stag2* in *Stag2^f/^*^y^ and *Stag2^f/y^;Cre* forebrains. Maximum cutoff of 3 was used.

**Figure S7.**
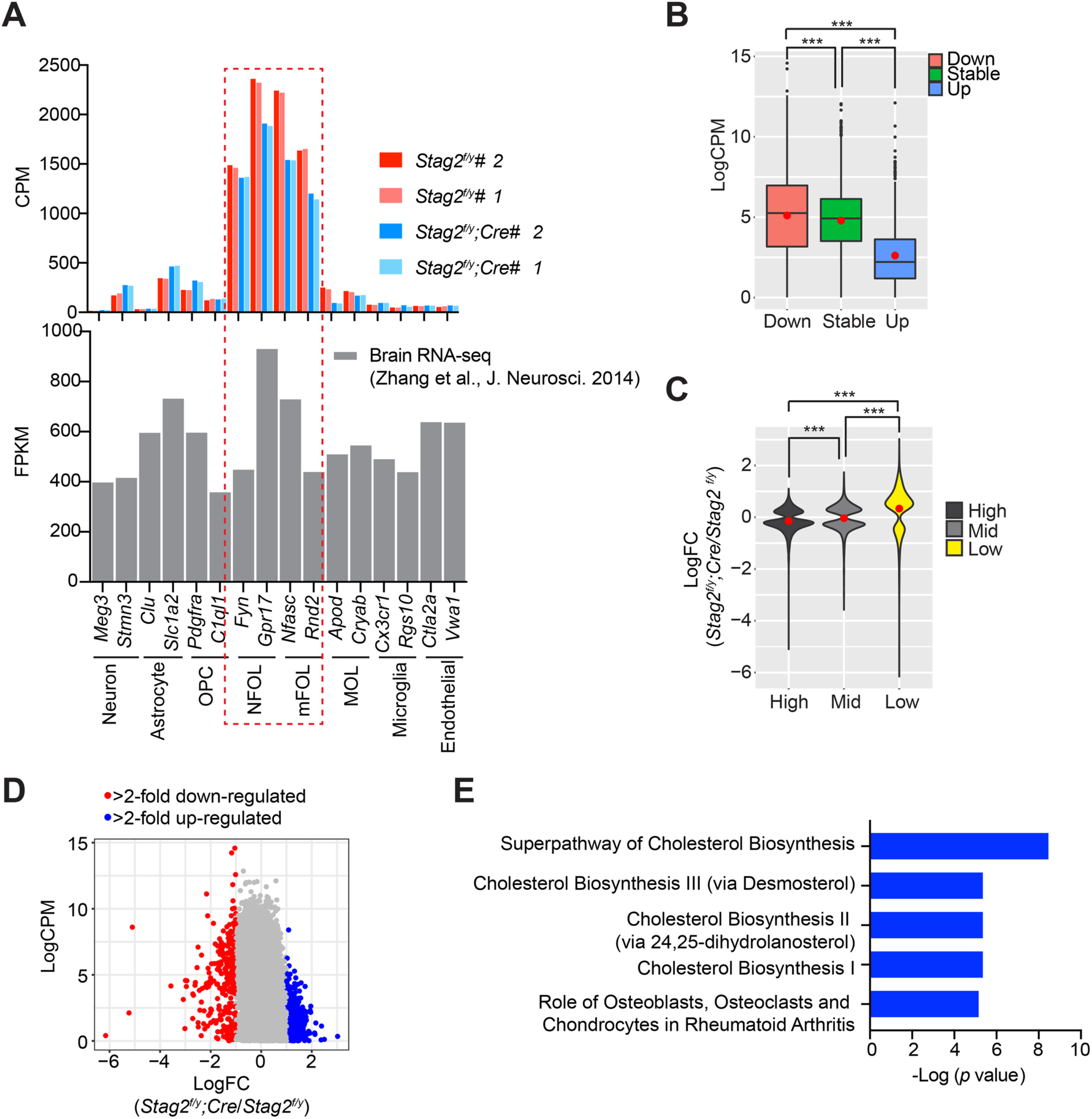
STAG2 regulates transcription in primary oligodendrocytes. (A) The expression levels of signature genes of indicated brain cell types in the isolated primary oligodendrocytes (OLs) in this study. The expression levels of the same set of signature genes in the individually isolated cell types from previous studies are shown below. NFOL and mFOL signature genes are highly enriched in the isolated primary OLs in this study. (B) Boxplot of the expression levels for genes in the indicated categories. Red dots represent the mean values. ***p < 0.001. (C) Violin plot of the expression changes for the active genes with different expression levels. Red dots represent the mean value. ***p < 0.001. (D) Scatter plot of the gene expression level against transcriptional changes. (E) IPA analysis of the down-regulated gene sets.

**Figure S8.**
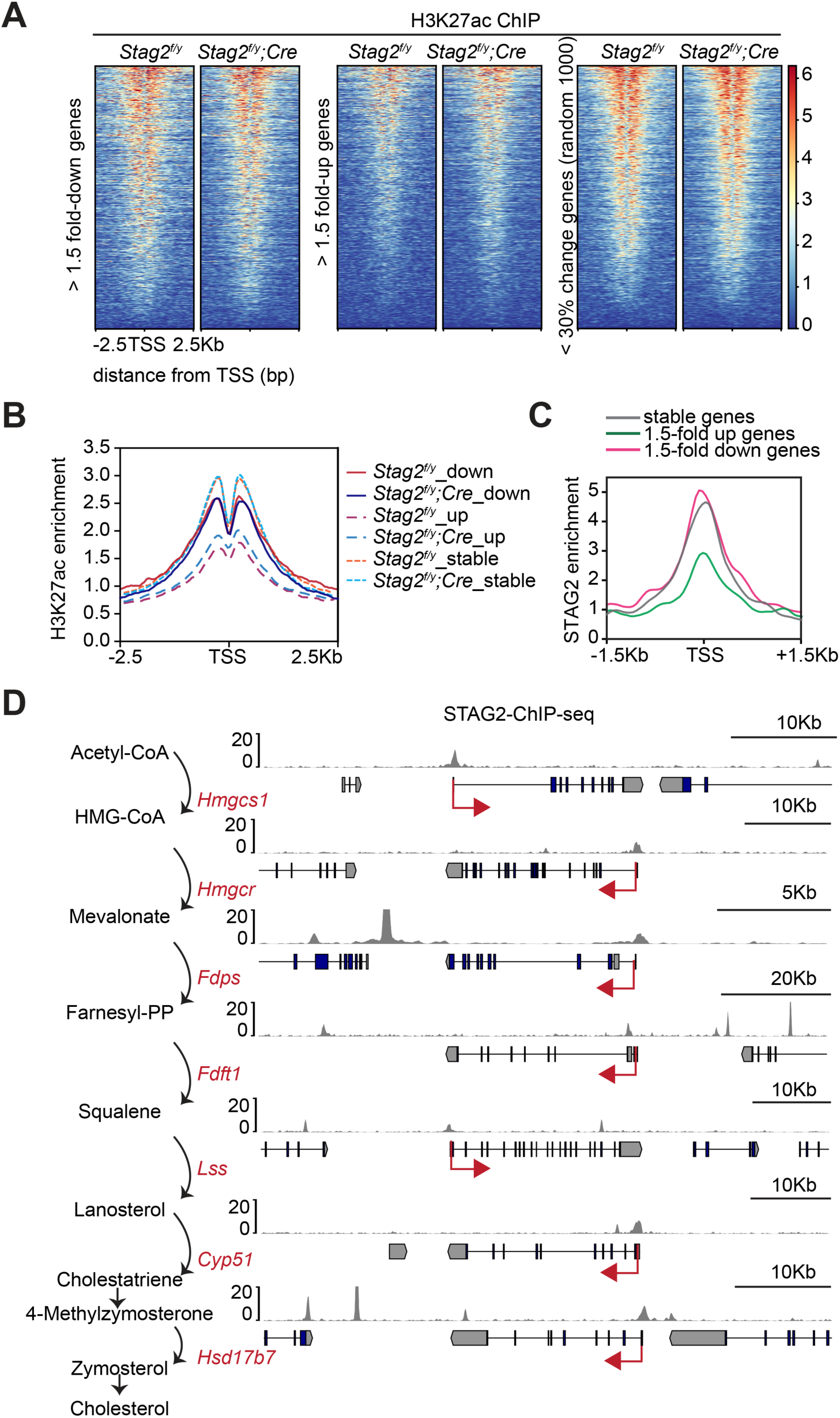
Enrichment of STAG2 and histone modifications at gene promoters. (A) Heatmap of H3K27ac ChIP-seq signal enrichment in the promoter regions of genes in the indicated categories. (B) Density profile of H3K27ac ChIP-seq signal enrichment in the promoter regions of genes in the indicated categories. (C) Density profile of STAG2 ChIP-seq signal enrichment in the promoter regions of genes in the indicated categories. (D) Binding of STAG2 at the genomic loci of down-regulated genes that encode cholesterol biosynthetic enzymes as revealed by ChIP-seq.

**Figure S9.**
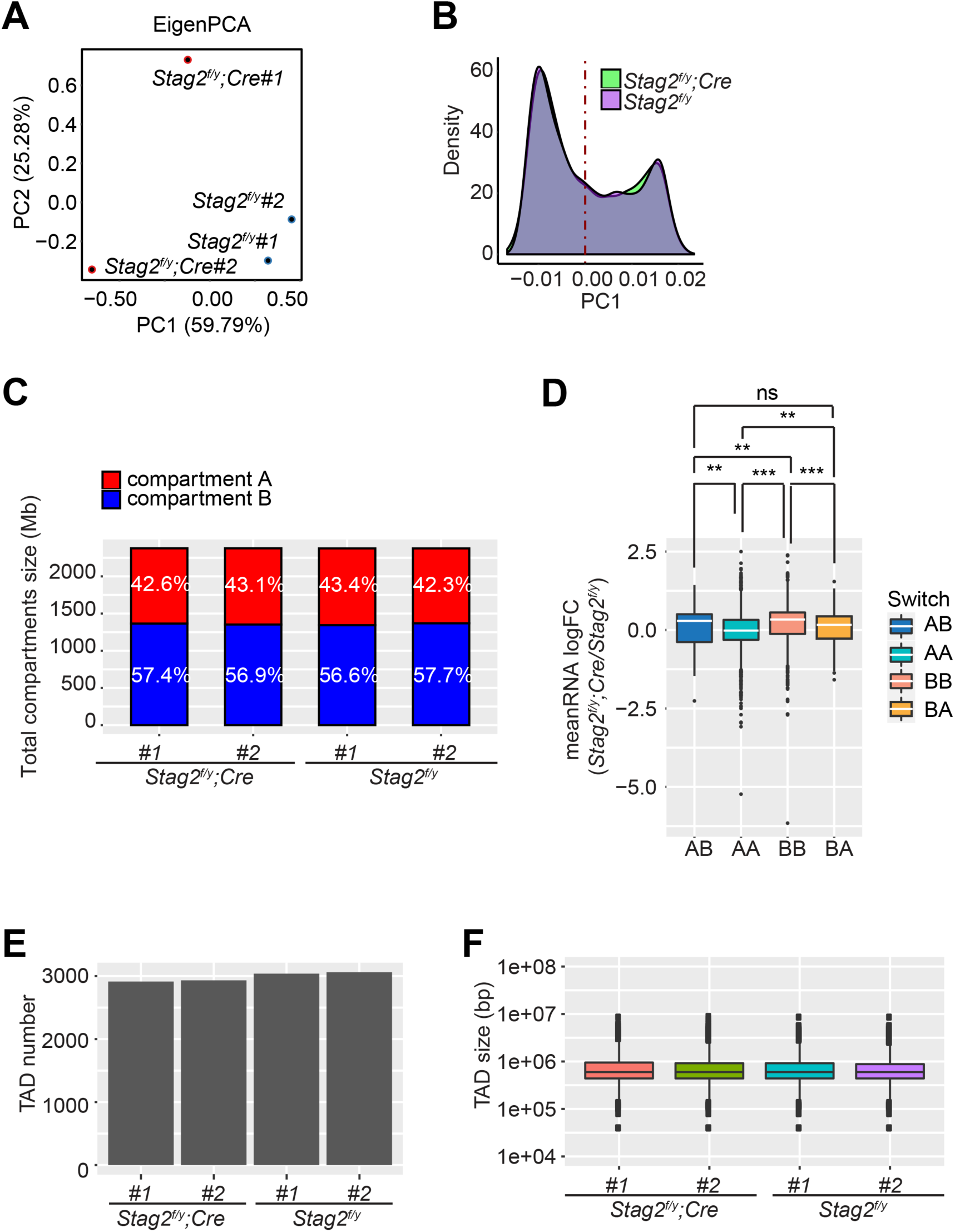
*Stag2* deletion does not alter chromatin compartmentalization or TAD formation in oligodendrocytes. (A) Principal component analysis (PCA) plot of the eigenvectors of the indicated samples. (B) Density plot of principal component 1 (PC1) of *Stag2^f/y^* and *Stag2^f/y^;Cre* oligodendrocytes. (C) Compartment compositions of the indicated samples. (D) Boxplot of the average gene expression change for all the differentially expressed genes (FDR < 0.05) inside each genomic bin. Bins counted: AA, 5806; AB, 155; BA, 251; BB, 2659. The unpaired Wilcoxon-test was used for the statistical analysis. *p < 0.05; **p < 0.01; ***p < 0.001; ns, not significant. (E) TAD numbers of the indicated samples. (F) TAD size distributions of the indicated samples.

**Figure S10.**
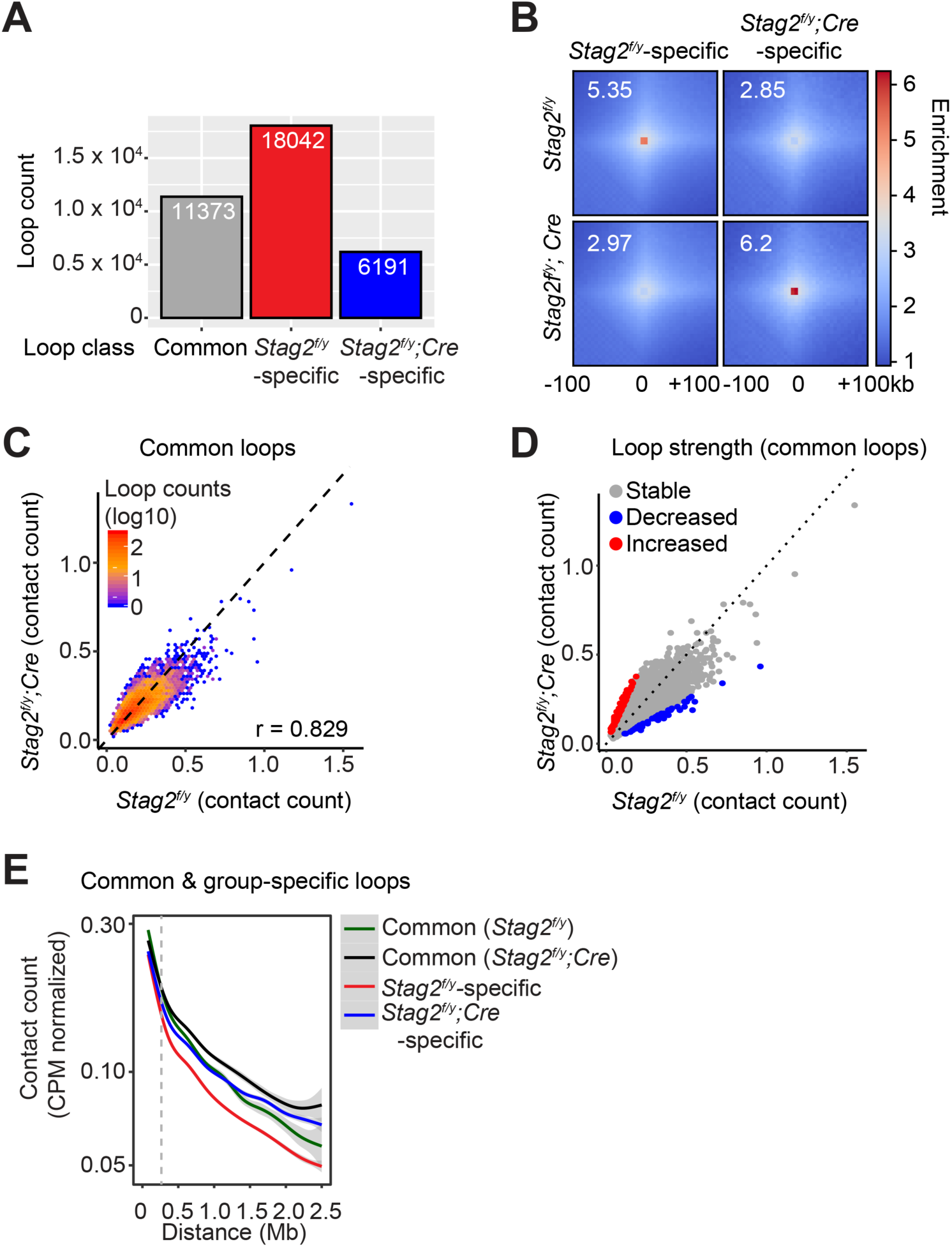
*Stag2* deletion reduces chromatin loops in oligodendrocytes. (A) Loop numbers of the indicated categories. (B) Pile-up analysis of loop “dots”-centered local contact maps for loops specific to *Stag2^f/y^* or *Stag2^f/y^;Cre* oligodendrocytes (OLs). (C) Hexbin plot of contact counts of common loops in *Stag2^f/y^* and *Stag2^f/y^;Cre* OLs. (D) Scatter plot of contact counts of common loops in *Stag2^f/y^* and *Stag2^f/y^;Cre* OLs. Loops with significantly changed strength in *Stag2^f/y^;Cre* OLs are highlighted in red (increased) and blue (decreased). Log_2_FC threshold of 1 was used. (E) Normalized contact counts for loops in the indicated categories across different genomic distances.

**Figure S11.**
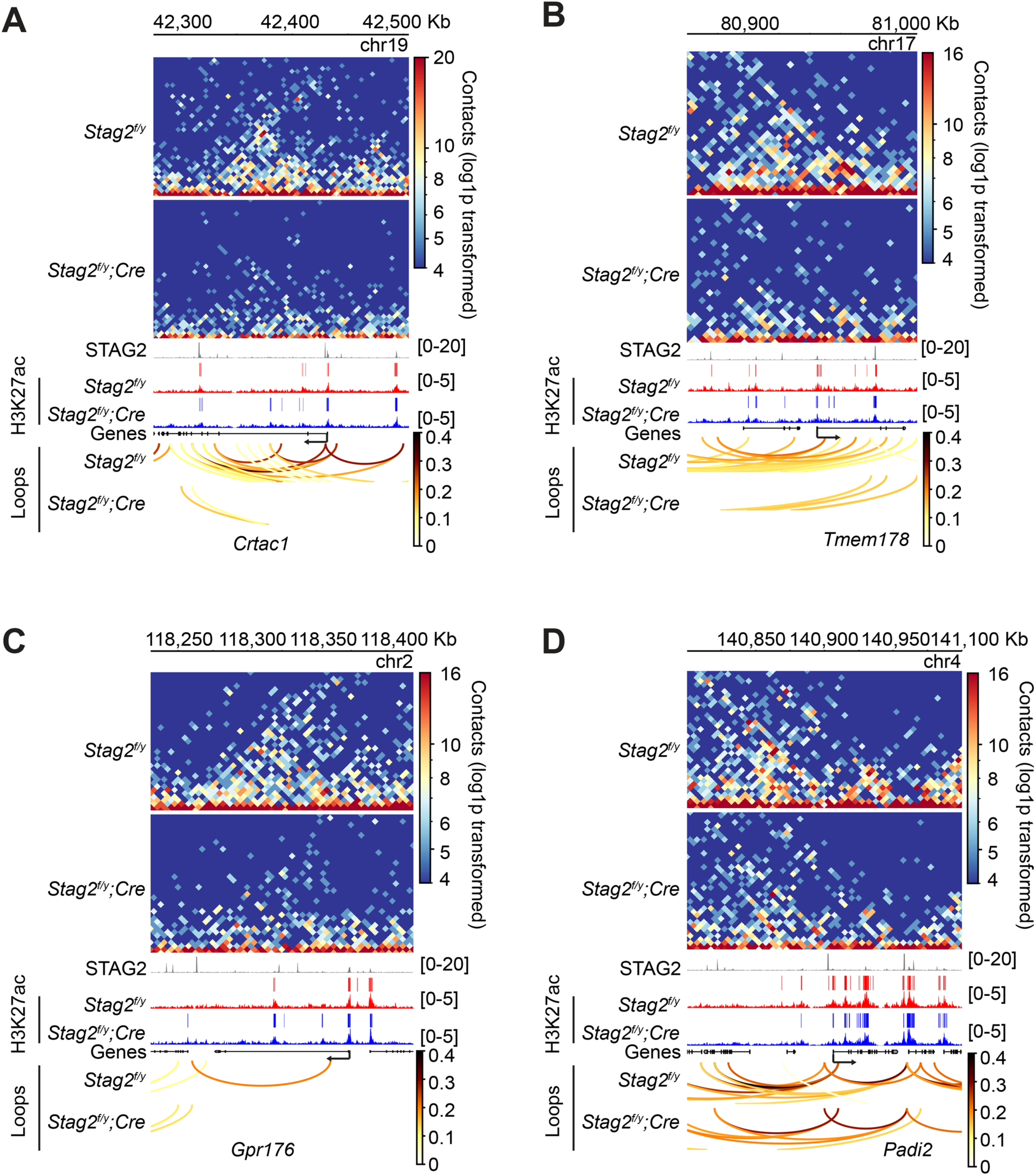
STAG2 controls local chromatin looping at differentially regulated genes. (A-D) Snapshots of the contact maps at the indicated differentially regulated genes. Tracks and peaks from STAG2 and H3K27ac ChIP-seq as well as loops are shown below. Transcription directions are indicated by arrows.

**Figure S12.**
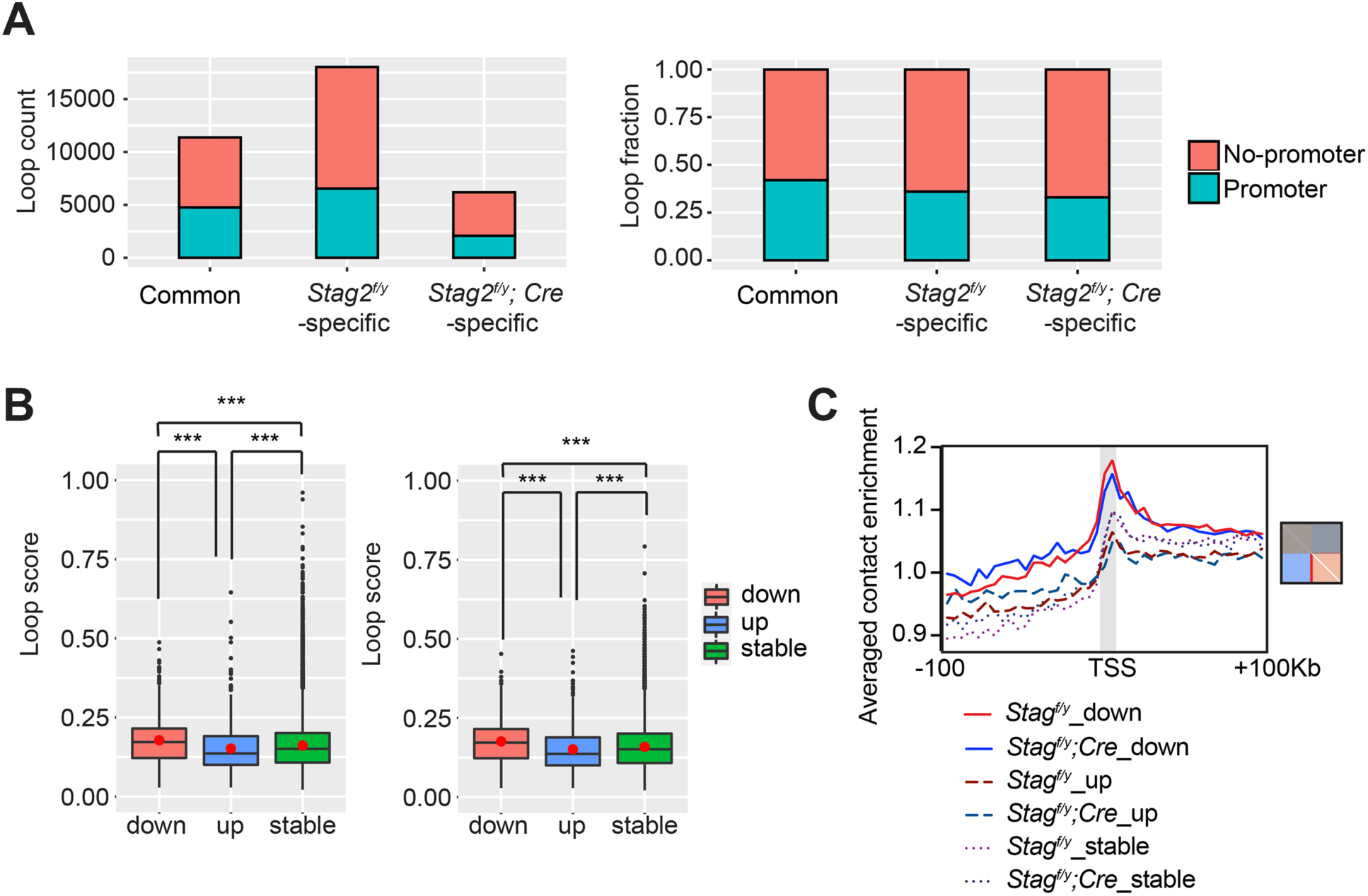
STAG2 regulates the formation of promoter-anchored loops in oligodendrocytes. (A) Loop counts (left panel) and fractions (right panel) of loops anchored at promoter or non-promoter regions in the indicated categories. (B) Loop score of the loops in Figure 6E. Loop score from *Stag2^f/y^* oligodendrocytes (OLs) was used for common loops on the left, and loop score from *Stag2^f/y^;Cre* OLs was used for common loops on the right. The unpaired Wilcoxon test was used for the statistical analysis. ***p < 0.001. (C) Profile plot of the average enrichment score for the bottom half of each graph panel in Figure 6F. Diagonal pixels were omitted.

## Notes

### Competing Interest Statement

The authors have declared no competing interest.

### Summary of Updates

Method details included.

